# Genotypic diversity and dynamic nomenclature of *Parechovirus A*

**DOI:** 10.1101/2020.08.14.251231

**Authors:** Edyth Parker, Alvin Han, Lieke Brouwer, Katja Wolthers, Kimberley Benschop, Colin A. Russell

**Author notes:** corresponding author: EP; CAR.

## Abstract

Human parechoviruses (PeV-A) can cause severe sepsis and neurological syndromes in neonates and children and are currently classified into 19 genotypes based on genetic divergence in the VP1 gene. However, the genotyping system has notable limitations including an arbitrary distance threshold and reliance on insufficiently robust phylogenetic reconstruction approaches leading to inconsistent genotype definitions. In order to improve the genotyping system, we investigated the molecular epidemiology of human parechoviruses, including the evolutionary history of the different PeV-A lineages as far as is possible. We found that PeV-A lineages suffer from severe substitution saturation in the VP1 gene which limit the inference of deep evolutionary timescales among the extant PeV-A and suggest that the degree of evolutionary divergence among current PeV-A lineages has been substantially underestimated, further confounding the current genotyping system. We propose an alternative nomenclature system based on robust, amino-acid level phylogenetic reconstruction and clustering with the PhyCLIP algorithm which delineates highly divergent currently designated genotypes more informatively. We also describe a dynamic nomenclature framework that combines PhyCLIP’s progressive clustering with phylogenetic placement for genotype assignment.

## Introduction

Human parechoviruses (PeV) of the species *Parechovirus A* of the *Picornaviridae* family (PeV-A) are globally prevalent pathogens that largely cause subclinical, mild respiratory or gastrointestinal disease in neonates and children. However, these viruses are also associated with more severe conditions including sepsis and neurological syndromes.^1^ The two prototypic strains of PeV-A were first isolated in the United States of America in 1956 from infants with diarrhoea.^2^ The two serologically distinct prototypes were initially classified as echovirus 22 and 23 in the *Enterovirus* genus, owing to similar clinical properties and cytopathology to other enteroviruses.^2^ Further investigation revealed that these viruses had distinctive molecular properties to enteroviruses, including high levels of sequence divergence as well as dissimilarities in genome structure and host cell protein interaction, and they were reclassified into a distinct *Picornaviridae* genus *Parechovirus* as the genotypes PeV-1 and PeV-2.^3^

To date, 19 genotypes have been proposed for PeV-As based on a 25% nucleotide sequence divergence threshold in the VP1 gene. The VP1 gene encodes the major structural protein of the icosahedral capsid. Notably, the VP1 protein of some PeV-A genotypes has an arginine-glycine-glutamic acid sequence (the canonical RGD motif found in several other picornaviruses) near the C terminus which mediates attachment to cell surface integrins.^1^ Genotypes PeV-3 and PeV-7 through 19 consistently lack the RGD motif in the VP1 gene and are presumed to be integrin-independent.^1^ The region encompassing the receptor binding site contains antigenic sites and is highly immunogenic.^4–6^

Of the 19 current PeV-A genotypes, PeV-1 and 3 are the predominant genotypes globally both in seroepidemiological studies and clinical settings, but prevalence of the individual genotypes vary widely across countries.^1,7–9^ PeV-1 is associated with mild gastrointestinal or respiratory symptoms, in children between 6 months and 5 years of age, whereas PeV-3 is more likely to cause severe disease in children under the age of 3 months.^7^ Central nervous system conditions such as acute flaccid paralysis, meningitis and encephalitis are more often associated with PeV-3.^10–13^ Inference about differential clinical manifestations is limited for most of the other PeV-A genotypes as they have only been isolated from a few cases.

The PeV-A genotyping criteria have changed over time.^14^ For example, the PeV-3 genotype was first classified based on an uncorrected nucleotide sequence distance to other PeV-A genotypes of more than 30% in the VP1 genomic region, a threshold that is also used for strain classification of enteroviruses.^15^ Several other thresholds on the nucleotide (23%, 27%) and amino acid level (13%, 19%) have been proposed for the VP1 gene as well as the VP3/VP1 junction (18% nt and 8% aa distance), but these thresholds have not uniformly applied.^15–17^

It is unclear if the current genetic distance based genotyping system delineates the population structure of PeVs at the appropriate resolution to capture the epidemiological and/or evolutionary processes underlying PeV-A diversity. The use of a distance threshold from a closely related pathogen to classify genotypes is a prevalent convention for less well-characterized viruses on discovery. This is typically because there is rarely any additional, systematic information on the epidemiology or serology of the virus to calibrate biologically meaningful limits on the genetic divergence allowed within demarcated groups. However, the assumption that information used to delineate thresholds in one virus may be generalizable to others is problematic. The high degree of amino acid sequence divergence in the capsid protein between PeV-1 and PeV-2 is comparable to the threshold delineating serotypes in better characterized picornaviruses such as enteroviruses and foot-and-mouth disease viruses.^1^ However, there has been no systematic characterization of the serological properties of PeVs, especially with the discovery of several novel, divergent lineages in recent years.^18,19^ It is therefore unclear whether genotypes characterized by high degrees of genetic heterogeneity in the VP1 region are capable of cross-neutralization. Cross neutralization has been reported between PeV1-2 and PeV4-6, with only PeV-3 being consistently antigenically unique. ^18,19^ The inconsistent information between antigenic and genetic diversity as currently described along with the lack of systematic antigenic characterization of PeVs suggest that antigenic information cannot be used to underlie the genotyping nomenclature PeV-A.

The current genotyping system also has other limitations. It operates on uncorrected genetic distances, which severely underestimates the true evolutionary distance between viruses in the presence of substitution saturation and high heterogeneity in lineage-specific evolutionary rates.^20,21^ Given the extent of genetic divergence among extant PeV-A, saturation is likely to be a substantial issue. The current genotyping system also relies on phylogenetic reconstruction approaches such as neighbor joining that can result in severe branch underestimation and topological inconsistencies across studies, which has resulted in inconsistent typing of PeV-A.^22–29^ It has also resulted in the designation of PeV-1 subtypes A and B, which do not consistently group as a clade.^30^ Many studies also employ BLAST-based nucleotide searches to compare query sequence to a reference dataset representing different genotypes.^22–28^ However, many of these reference sequences are not representative of the high internal divergence of some genotypes. This is especially pronounced for the progenitor sequences of some genotypes such as PeV-1 and 3, which are genetically and antigenically distinct.^22–28^ Inconsistently typed viruses are incorrectly annotated in public sequence databases, propagating the error into subsequent studies.

The inconsistent discriminatory information in the presently designated genotypes highlights fundamental limitations of the current genotyping system. Nomenclature systems should aim to delineate populations into units of genotypic similarity that carry cohesive and consistent information about the evolutionary dynamics of pathogen variants that result in differences in epidemiological and virologic characteristics. Here, we describe and quantify the evolutionary history of the different PeV-A lineages as far as is possible with currently available data and tools and propose an alternative, dynamic PeV-A nomenclature system based on robust phylogenetic reconstruction methods and clustering with PhyCLIP in combination with a phylogenetic placement algorithm for the assignment of genotypes to newly sequenced viruses.

Though demarcations of genotypes or lineages are often vague and boundaryless, the phylogenetic clustering algorithm PhyCLIP offers a statically principled and phylogenetically informed framework to partition phylogenies into discontinuous clusters that may represent independent evolutionary and epidemiological phylogenetic units.^31^

## Methods

### Dataset curation

All available nucleotide sequences (as of 01/06/2019) of the VP1 gene (±700 nt) with known dates of isolation were collated from Genbank (N=1655; i.e. the complete undownsampled dataset). The sequences were classified according to genotype identity annotated in the Genbank metadata or the associated literature. Unannotated sequences were classified as “unknown”. Sequences obtained from a recent Malawian cohort study (n=123) were excluded from this complete dataset, as they were reserved as a test dataset to assess the validity of the proposed dynamic nomenclature system (see phylogenetic placement section below and Results).^22^ Genbank accession numbers available at https://github.com/AMC-LAEB/parecho.

The sequences in the complete dataset are not uniformly distributed across the current genotypes, with representation skewed towards PeV-1 (n=608) and PeV-3 (n=614) (SFigure 1). To account for sampling biases, we downsampled PeV-1 and PeV-3 viruses from the complete dataset to the equivalent number of the next largest genotype (PeV-4: n=115; final included dataset of PeV-1: 117; PeV-3: 120) in the complete dataset to generate a *primary dataset* that was used for all subsequent analyses other than the molecular dating analyses. To obtain the primary dataset, we downsampled PeV-1 and PeV-3 viruses from the complete dataset to the equivalent number of the next largest genotype (PeV-4: n=115; final included dataset of PeV-1: 117; PeV-3: 120). Sequences were sampled at random but were drawn to maintain the genetic distance distribution of the complete dataset (Figure 1). We also removed three sequences (GenBank accessions: KM407606, KY931551 and KM407607) that were highly divergent (inferred branch length >0.4 substitutions/site) based on a preliminary maximum-likelihood phylogeny reconstructed under the GTRGAMMA substitution model in RAxML from a MAFFT-derived codon alignment of the primary dataset.^32,33^

**Figure 1:**
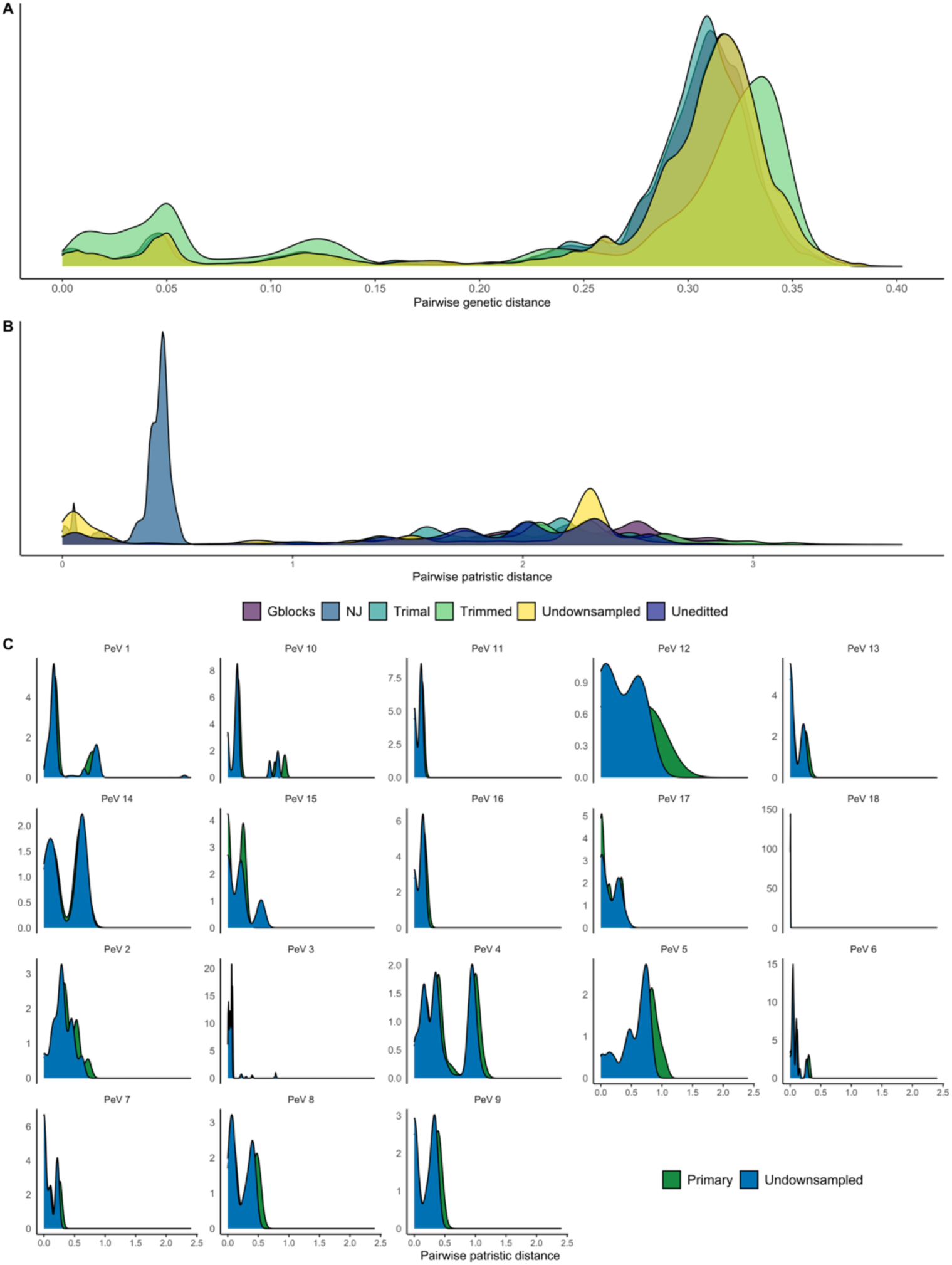
A) Density of pairwise genetic distance distribution in the nucleotide alignments. All alignments are of the primary dataset (green), except for the full undownsampled complete dataset (in yellow). B) Density of pairwise patristic distance distribution in the reconstructed nucleotide phylogenies. C) Within-genotype pairwise patristic distance, as designated by the current nomenclature in the primary and undownsampled dataset.

A sub-dataset consisting of all PeVs isolated before 2006 (referred to as the pre-2006 dataset) was used to assess the phylogenetic clustering algorithm PhyCLIP’s sensitivity to sampling (see Supplementary Information). Finally, a set of all available whole genome sequences (n=158) was collated from Genbank for recombination analysis.

### Sequence alignment and phylogenetic reconstruction

PeV-As are characterized by a high degree of genetic heterogeneity (Figure 1). The high level of sequence divergence in the VP1 region, particularly the C terminal region, (mean genetic distance of 28%) can result in ambiguously aligned regions, rendering the alignment unreliable and introducing systematic error into phylogenetic reconstruction and downstream phylogenetic-based genotyping inferences. However, removing potentially unreliable regions of the alignment results in a loss of information that could decrease phylogenetic signal substantially, which is of particular concern when working with very short subgenomic regions such as VP1 (~700nt). We constructed four different nucleotide alignments subject to various quality filtering specifications from the primary dataset, as well as an amino acid alignment, to investigate the robustness of phylogenetic inference in the trade-off between the potential loss of phylogenetic information and bias introduced by potentially misaligned regions or other model misspecifications:

1. A nucleotide alignment with no additional editing
2. A nucleotide alignment with the last 70 nucleotide positions encompassing the hypervariable, often ambiguously aligned RGD-motif region removed
3. A nucleotide alignment filtered to sites that passed TrimAL’s heuristic selection of quality control parameters based on similarity statistics.^34^
4. A nucleotide alignment filtered to sites extracted using GBlocks, allowing small blocks and gaps (b4=2 and b5=all).^35^
5. An amino-acid alignment, translated from the unedited alignment (alignment 1)

All nucleotide alignments were constructed using the codon model in PRANK and manually edited. PRANK is a phylogeny-aware alignment algorithm that employs ancestral reconstruction and has shown improved performances on alignments with insertions and deletions.^36^

We performed model adequacy tests to prevent substitution model underparameterization and model assumption violation, which are the primary sources of bias in phylogenetic reconstruction.^37^ Potential sources of error include substitution saturation and model misspecification regarding stationarity, rate variation across branches and sites as well as base composition heterogeneity. We used the chi-squared test implemented in IQ-TREE to investigate heterogenous base composition among lineages, which would violate model assumptions of stationarity.^37,38^ We performed model fit tests to identify the substitution model with the highest statistical fit by Bayesian Information Criterion (BIC) for each alignment within IQTree (Table 1).^39^ Phylogenetic trees were reconstructed for each of the alignments in IQTree under the best performing models, with 1000 ultrafast bootstrap approximation (UFBoot) replicates employing the BNNI hill-climbing nearest neighbour interchange search for further optimization of each bootstrap tree to reduce the risk of nodal support overestimation.^40^ Additionally, a phylogenetic tree was reconstructed with Neighbour Joining under the Kimura 2-parameter substitution model in MEGA to investigate the systematic underestimation of branch lengths by more simple phylogenetic approaches.^41^ Phylogenies constructed from the different alignments were compared with tanglegrams produced with the Baltic module (https://github.com/evogytis/baltic). All calculations of phylogenetic statistics, including patristic distance, were performed with the *ape* package in R.^42^

**Table 1:**
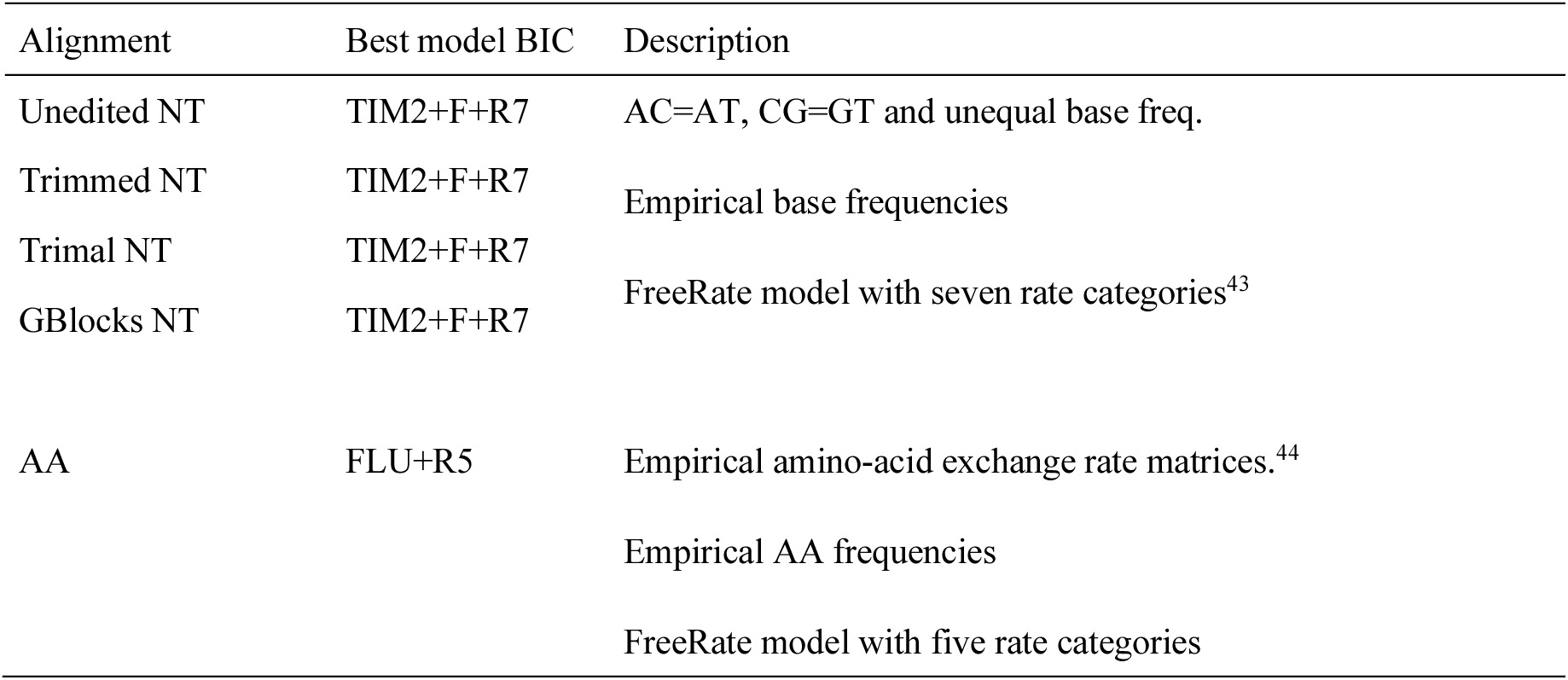
Model fit for phylogenetic reconstruction of the primary dataset

Due to the high degree of genetic divergence among PeV-A genotypes, we used three approaches to evaluate the extent of mutation saturation in the nucleotide alignments. Substitution saturation occurs when there are multiple unobserved substitutions at a single site which are not accounted for when modelling sequence divergence to branch lengths, resulting in systematic underestimation of the branches in the tree.^45^ Saturation mostly occurs at the rapidly evolving third codon position of nucleotide sequences where there is a high probability of synonymous substitution. Standard evolutionary models that do not account for variation of selective pressure and the associated rate heterogeneity across sites and branches are known to significantly underestimate branch lengths, especially branches under strong purifying selection.^20^ However, conventional model fit tests applied in phylogenetic reconstruction do not assess how models accounts for substitution saturation, and do not allow for the rejection of all models if all models are poor descriptions of the evolutionary process that generated the data. ^37,45^

First, transition and transversion frequencies were plotted against genetic distance to visualise the extent of saturation. Second, we formally tested for saturation with the information entropy-based index of substitution saturation using the Xia’s test as implemented in DAMBE.^46^ The alignment was partitioned into a combination of the first and second codon site and the third codon site separately, to account the different rates of synonymous and non-synonymous substitutions and saturation at the different sites. In the third approach, the branch lengths for the maximum-likelihood phylogenies constructed from the different nucleotide alignments were re-estimated in HyPhy under two different models of evolution: the standard GTR substitution model with gamma-distributed site rate variation across four categories and the selection-aware aBSREL model.^47^ aBSREL is a branch-site random effects likelihood (BSREL) model that estimates the effects of varying selection pressure across codon sites and branches by inferring dN/dS (non-synonymous or non-silent to synonymous or silent substitution rate ratio) rate classes for each branch and estimating the proportion of sites evolving under each rate class.^47^

### Selection analysis

Selection analysis was performed with a set of codon models implemented in HyPhy.^48^ Fast, Unconstrained Bayesian AppRoximation (FUBAR) was used to detect pervasive positive or purifying selection at individual sites.^49^ Site-specific selection was investigated with the Mixed Effects Model of Evolution (MEME).^50^ MEME is currently the most robust site-to-site selection approach and is more comprehensive than FUBAR as it accounts for both pervasive and episodic selection.^50^ As above, the adaptive Branch-Site Random Effects Likelihood (aBSREL) was used as a branch-site model of selection, allowing evolutionary rates to vary across lineages and sites to detect lineage-specific positive diversifying selection.^47^

### Recombination analysis

Alignments of the primary dataset were screened for recombinant sequences with the RDP, GENECONV, MAXCHI, CHIMAERA, 3SEQ, BOOTSCAN and SISCAN tests implemented in the RDP program suite.^51^ Default settings were used, excluding the window size, which was set to 30bp across tests. Potential recombinants were defined as those with Bonferonni corrected p-values below 0.05 in more than three detection methods. The full genome dataset was screened for recombination with the Genetic Algorithm for Recombination Detection GARD, implemented in Hyphy.^52^

### Evolutionary history of PeV

We wanted to investigate the evolutionary timescale and relationship of the divergent PeV-A genotypes, with particular interest in dating the divergence between the co-circulating genotypes. Divergence dating requires that the genetic divergence of sequences be scaled to units of absolute time by assuming a molecular clock model, informed by tip-calibrations.^53^ Molecular clock models are a statistical description of the relationship between observed genetic distances and time. In other words, prior to divergence dating, we must ensure that there is sufficient temporal signal in the dataset to inform clock models.^54^ The strength of the temporal signal required for molecular clock analyses was investigated in three different datasets:

1. The complete, non-downsampled dataset
2. The primary dataset downsampled to maintain extant diversity, in both amino acid and nucleotide reconstructed phylogenies
3. The individual sublineages of PeVs, here defined by the existing PeV-A genotyping system prevalently used.

Phylogenetic trees were reconstructed for each of the sub-lineage datasets with RAxML v8.2.1.1 under the GTR+gamma4 model.^32^ The temporal information for each dataset was quantified with root-to-tip regression, performed in R with the *ape* package.^42^ Calibration was based on sampling times, resolved to the year. Clocklike structure was evaluated for hypothetical roots including midpoint rooting and root positions placed to maximize the correlation between tip sampling dates and distance to root and minimize the sum of the squared residuals in the regression.

The evolutionary rate and history of the individual PeVs genotypes was estimated using Bayesian Evolutionary Sampling Trees (BEAST) software package version 2.5.1.^55^ The alignments were partitioned into codon positions 1+2 and 3 to account for variation in rates across sites, with a GTR gamma model with 4 rate categories as substitution model in respective partitions but shared clock and tree models. A lognormal relaxed clock was employed to account for the high variation in rates across branches suggested in temporal regression. A Bayesian skyline and constant size coalescent model were used as tree model priors respectively (See Supplementary files). Chains were run for 500 million steps across the datasets (Stable 7), with convergence diagnosed as an estimated sample size in all parameters > 200 in Tracer v 1.7.1.^56^ Log- and tree-files from individual runs were combined and sub-sampled with LogCombiner were necessary. All runs were also completed by sampling from the prior as diagnoses.

### Phylogenetic clustering with PhyCLIP to define genotypes

Phylogenetic clustering was performed on the phylogenies constructed from the primary, pre-2006 and primary with Malawian test sequence datasets using PhyCLIP.^31^ PhyCLIP operates on the distribution of all branch lengths in the phylogeny, using this global patristic distance distribution as a pseudo-null distribution to test the within-cluster distance distribution of putative clusters against. PhyCLIP also incorporates the branching order of the phylogeny into cluster definition through its distal dissociation approach, which accommodates the designation of paraphyletic clusters and performs outlier testing. PhyCLIP was run with different sets of the parameters varying over the ranges: a minimum cluster size of 2–10, a multiple of deviation (γ) of 1–3, and an FDR of 0.05, 0.1, 0.15, or 0.2. The optimization criteria were ranked as 1) percentage of sequences clustered, 2) grand mean of within-cluster patristic distance distribution, 3) mean of the intercluster distances. Percentage sequences clustered was prioritized as optimization criteria to assign the maximum number of sequences to clusters. Mean within-cluster distance was minimized to ensure clusters of closely related sequences were recovered, while inter-cluster distance was maximized to ensure well-separated clusters.^31^

### Phylogenetic placement to rapidly genotype new viruses

While PhyCLIP can be used to delineate diversity into statistically supported units and progressively update the nomenclature system when additional PeV-A diversity is sampled, rerunning the entire phylogenetic and PhyCLIP to genotype new individual or small numbers of viruses is very time consuming. Alternatively, phylogenetic placement can be used to rapidly genotype newly sampled viruses to mitigate the need to conduct full phylogenetic reconstruction and PhyCLIP clustering analyses for every new query sequence.

We employed a leave-out testing approach to validate phylogenetic placement on the Malawian cohort test dataset that included 123 viruses from a Malawian cohort.^22^ The RAxML-EPA PROTGAMMAGTR substitution model was used to phylogenetically place the Malawian sequences on the primary phylogenetic tree based on amino acid sequence alignment. The RAxML-EPA approach was originally developed for rapid phylogenetic classification of short-read sequences obtained from metagenomic studies but can also be applied to longer sequences. The algorithm traverses along all edges of the reference phylogenetic tree and computes a tree likelihood score under the maximum-likelihood model as it inserts the query sequence along each edge. The query sequence is phylogenetically placed on the best scoring edge and the corresponding normalized likelihood score (i.e. likelihood weight ratio, LWR) can be used as a measure of placement uncertainty.^57^ Under our framework (Figure 2), a query sequence would be typed to a cluster if it is topologically placed on an edge that was subtended within the cluster in the reference phylogeny and if the length of the inserted query branch estimated by RAxML-EPA does not exceed the maximum within-cluster pairwise divergence across all clusters. A cluster wide LWR is also computed by summing the LWR score of all edges within the cluster.

**Figure 2:**
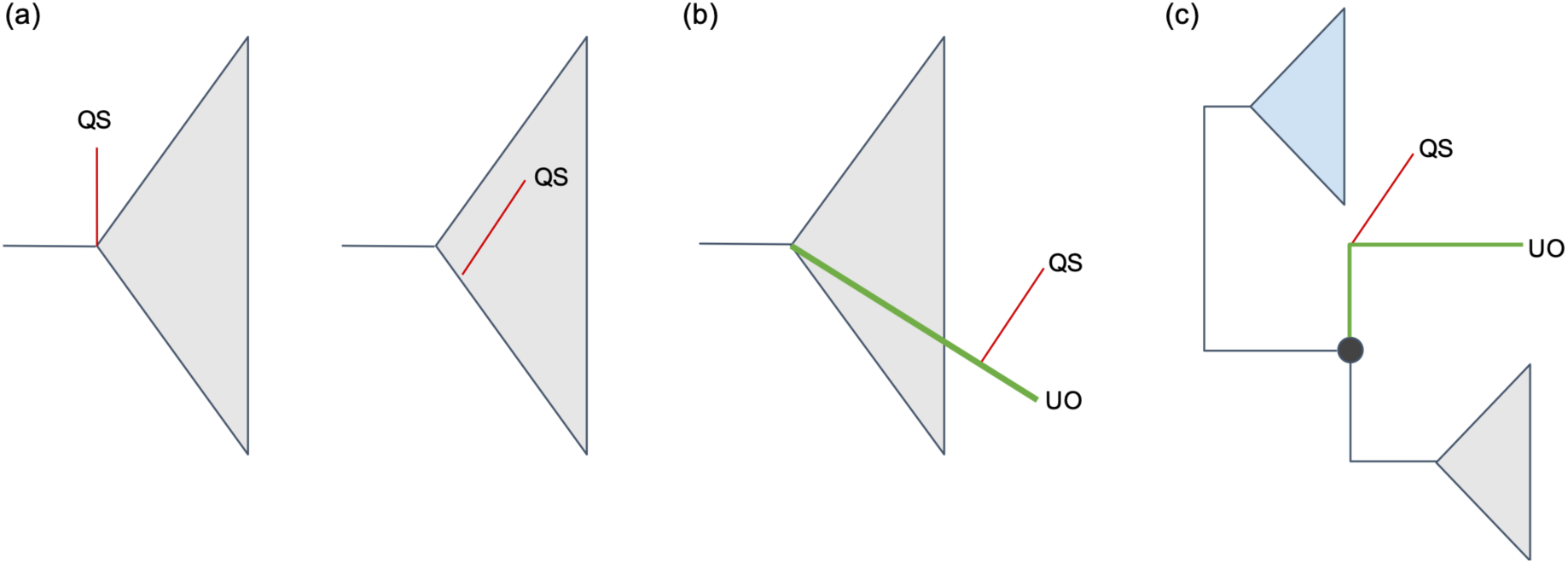
Types of query sequence (QS) based on its phylogenetic placement relative to PhyCLIP-defined clusters. (a) QS is closely related to reference viruses clustered as a single phylogenetic unit by PhyCLIP (grey triangle). (b) QS is placed on an unclustered outlying (UO) sequence relative to a PhyCLIP cluster (grey triangle). (c) QS is placed on any UO sequence or lineage that topologically lies between different PhyCLIP clusters.

To assess the accuracy of the phylogenetic placement as well as consistency of clustering topology between the reference and test phylogenies, a phylogenetic tree was reconstructed from the amino acid alignment of the test data with the additional Malawian sequences in IQTree under the best performing model selected by BIC, with 1000 ultrafast bootstrap replicates and bootstrap tree optimization with the BNNI algorithm.^38^ The phylogenetic placement of each query sequence was compared to its position in the reconstructed phylogeny. Code and data are available at https://github.com/AMC-LAEB/parecho

## Results

### Alignment quality and model adequacy in phylogenetic reconstruction of highly divergent viruses

All alignments showed extensive genetic divergence among genotypes and even within some genotypes (Figure 1). For each of the four nucleotide and one amino acid alignments, chi-square tests did not find evidence that any sequence or lineage significantly deviated in base composition from the dataset average. All of the substitution models ranked as best-performing by the Bayesian information Criterion (Table 1) included the FreeRate model of rate heterogeneity across sites with seven rate categories, as this extremely flexible site-to-site variation improves the accuracy of branch length estimation.^20,43,58^

A robust root is required to interpret directionality of evolutionary events in phylogenies and for the reconstruction of ancestral states. Well-resolved rooting of the PeV-A phylogeny was problematic, as conventional approaches showed severe violations of necessary assumptions. The phylogeny could not be rooted by temporal structure as there was no temporal signal in the dataset (see ‘Evolutionary History’ section below). There was also no clear outgroup to the full phylogeny, as large evolutionary distances to its closest potential outgroup virus, the Llungan parecho B virus isolated from bank voles, risks the introduction of rooting artefacts.^59,60^ This also extends to rooting to precursors of specific genotypes, such as the Harris strain. Long branch attraction from large evolutionary distances can be overcome by approaches limiting substitution saturation, including the exclusion of the rapidly evolving third-codon site, but the potential loss of information from the already short VP1 subgenomic region precluded this option.^61^ We opted for midpoint rooting, which assumes that all lineages evolve at the same rate. This assumption is highly likely to be violated in the dataset (see following section) but was chosen as the least problematic option.

The phylogenies reconstructed from all four nucleotide alignments were characterized by deeply divergent lineages, separating major clades with long interior branches (Figure 3A). The branching order of the major clades across the phylogenies reconstructed from the different nucleotide alignments were similar, although branch lengths varied (Sfigure 2). One exception was that Genotype 13 was incongruently placed between the phylogenies constructed from the unedited/Gblocks alignment and the trimmed/Trimal alignments. In the unedited/Gblocks phylogenies genotype 13 is basal to the subtree encompassing genotypes 1, 6, 16 and 18, whereas it is placed as a sister clade to genotypes 8 and 9 in the trimmed/Trimal reconstructed alignment. Both these bifurcations have low nodal support (40-44) across the trees. Notably, the phylogenetic trees reconstructed from the unedited alignment, which retains the most information, and GBlocks alignment, which had the most conservative filter for alignment quality, had identical topologies. Resultantly, we focused on the unedited alignment in the subsequent section as it retains the variable C terminus but yielded phylogenetic topologies that were consistent with more conservative approaches.

**Figure 3:**
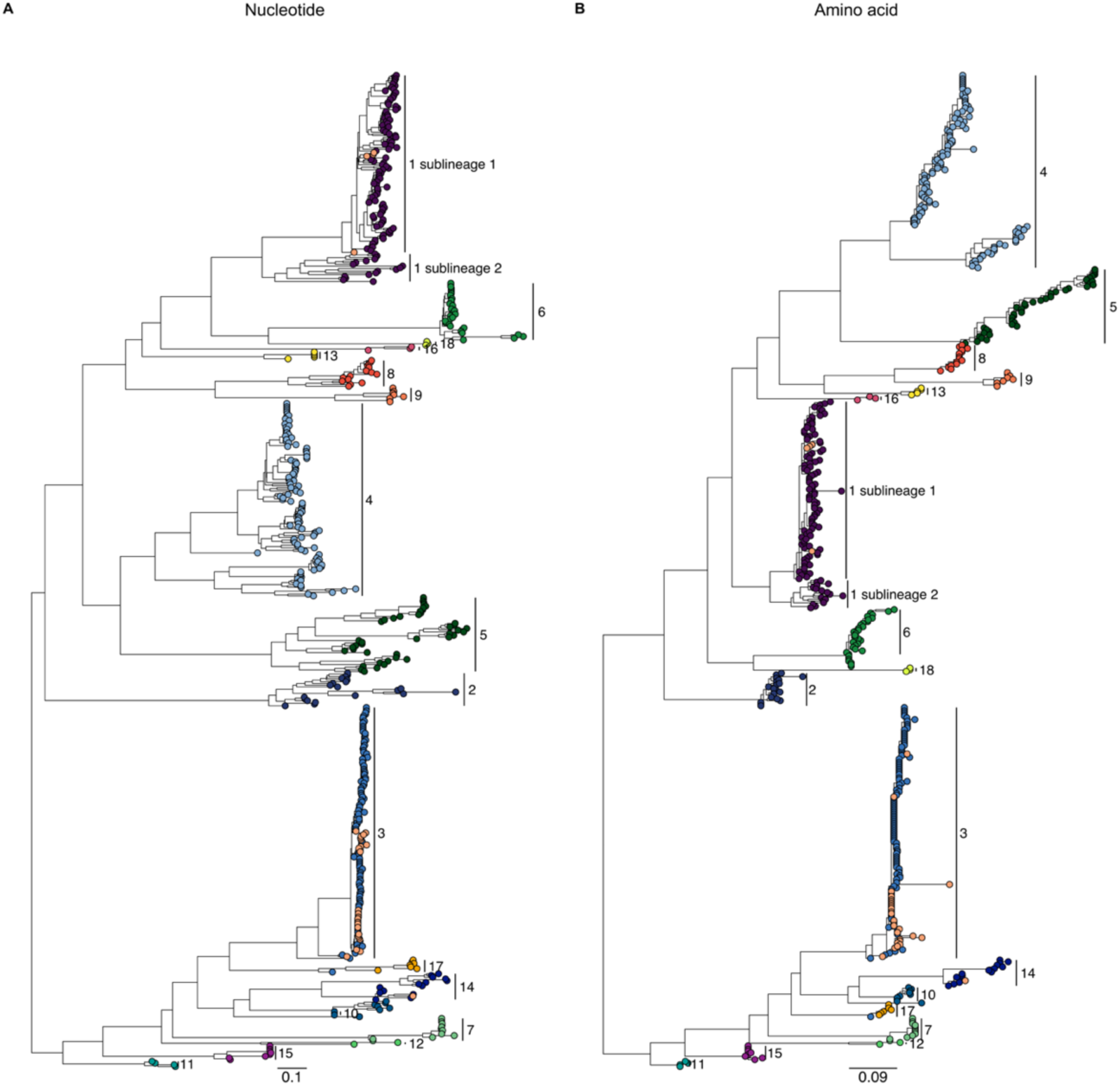
PeV-A phylogeny reconstructed from A) unedited nucleotide and B) amino acid alignment of the primary dataset. Tips colored by genotype defined under current system with associated labels. Light orange tips indicate sequences unclassified in Genbank metadata.

The number of long internal branches in the phylogeny (Figure 3A) strongly suggests potential biases in branch length estimation, including the possibility that phylogenetic signal has been confounded by substantial substitution saturation. We partitioned the alignment by codon positions (positions 1+2 and position 3 by itself) and found strong evidence for saturation at the third position using Xia’s test (p=0.001, see Methods). Phylogenetic analysis can be restricted to the first and second position to limit confounding by saturation.^62^ However, we excluded this option as phylogenetic information was already limited by the short length of the VP1 subgenomic region.

The systematic underestimation of branch lengths by conventional substitution models was investigated with branch re-estimation under the aBSREL model in Hyphy, following previous methods.^20^ There was strong evidence for variation in selection pressure across sites and branches over time, with the adaptive branch site-model that infers the optimal number of omega rate categories per branch showing the best fit by AICc (STable 1). The majority of branches in the tree could be sufficiently modelled with all sites evolving at a single rate, with a small proportion of branches (8.8%) best described with site-to-site variation modelled as two rate categories (STable 2). This includes the deepest branch defining two major clades: one encompassing genotypes 1, 6, 8, 9, 13, 16 and 18, which is under strong purifying selection (dN/dS = 0) at 96% of its sites, and incredibly strong diversifying pressure (dN/dS > 500) at the rest, the other encompassing genotypes 3, 7, 11-12, 14 and 17 (dN/dS =0, 92%, dN/dS > 200, 8% (STable 2).

There was no evidence for episodic diversifying selection in the full phylogeny, after correction for multiple testing across all branches. Exploratory testing of all branches for positive selection under aBSREL substantially reduces the statistical power of the test, particularly after conservative multiple test correction.^47^ Under the less conservative Benjamini-Hochberg FDR correction, seven branches approached significance (q<0.08), though interpretation is limited by the lack of statistical power owing to the penalty of multiple testing. This included two terminal branches, which is likely a result of model overfitting, the two long interior branches segregating the major clades in the deep topology of the tree, and branches that belonged to PeV-10 and 15 (STable 2). Selection analysis with FUBAR found no statistically significant evidence of pervasive diversifying selection, i.e. selective pressure aggregated over all branches, with posterior probability of 0.9.^49^ Episodic positive selection was detected at several individual sites by MEME (STable 4) when selection was performed for each genotype with sufficient samples (1-6) individually (p<0.05), though statistical power was limited.^50^ Signals for positive selection were found in the structured C-terminus as well as in a region that forms part of an epitope extending across subunits. However, detailed structural analyses of the capsid and its interactions have only been undertaken for PeV-1 and 3 thus limiting inferences about what phenotypes might be subject to selection.^63–65^

Branch length estimates were reasonably congruent between models for shorter branches (≤0.05 expected substitutions per site) across all datasets (Figure 4). The long internal branches were systematically underestimated by the GTR model compared to estimates under aBSREL. The expected number of substitutions per site along these branches suggest severe saturation, with branch length estimates approaching numerical infinity (Figure 4A).^58^ This includes the branches defining PeV-6, 13, 16 as well as the internal branches separating two major clades of PeV-5. Terminal branches estimated to have infinite lengths only have non-synonymous substitutions, resulting in an infinite omega parameter. Notably, these point estimates are likely to have extremely wide confidence limits, as modelling severe saturation can be imprecise.^20^ However, the result strongly supports that the depth of the PeV-A phylogeny is substantially underestimated by conventional substitution models.

**Figure 4:**
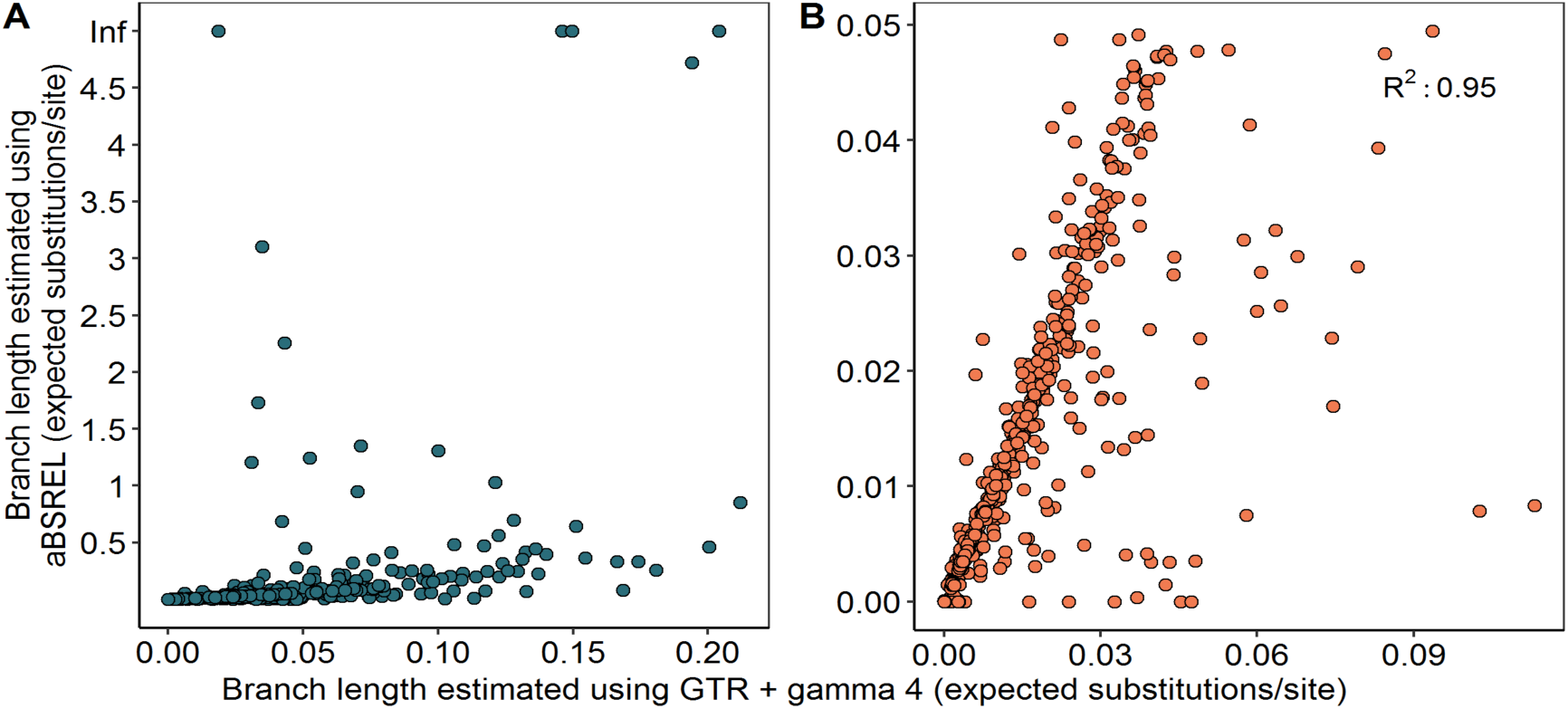
Branch length re-estimation under the aBSREL model. Branch lengths > 5 expected substitutions per site were coded as infinity for visualisation. A) All branches in the phylogeny. B) Short branches (<0.05 substitutions/site)

It is clear that numerous branches in the nucleotide-level phylogeny of PeV-A cannot be estimated with confidence due to the systematic underestimation of deep branches.^20^ This is likely to lead to systematic biases in phylogenetic reconstruction and subsequent inference of relatedness. This is particularly problematic for PeV-A as the current nomenclature system is predicated on reliable inference of extent of nucleotide divergence among genotypes. It is therefore more reliable to reconstruct PeV-A phylogenies from amino acid distances, which have a lower rate of evolution and are less likely to be affected by saturation.^37^ Phylogenies reconstructed from amino acid sequences are therefore more likely to accurately resolve the true evolutionary relationship of PeVs, even with the associated loss of information moving from nucleotide to amino acid data.

### Phylogenetic structure and diversity of PeVs partitioned by the current nomenclature system

As observed in the nucleotide-level phylogeny (Figure 3A), the amino-acid level phylogeny of the VP1 gene is characterized by long interior branches segregating major clades (Figure 3B). The two major clades stemming from the root bifurcation encompass currently defined PeV-A genotypes 1, 2, 4, 5, 6, 8, 9, 13, 16 and 18 and genotypes 3, 7, 10, 11, 12, 14, 15 and 17 respectively. The two distinct lineages are not differentiated by the presence of the RGD motif, as the second distinctive lineage (3, 7, 10, 11, 12, 14, 15 and 17) all lack the RGD motif, but as do genotypes 8, 9, 13, 16 and 18. The nodes defining the currently designated genotypes and the deeper bifurcations in the tree are supported by high ultrafast bootstrap approximation (aBS) bifurcation support values (aBS >0.7), though the bifurcation of PeV-1 from the clade encompassing current genotypes 4, 5, 9, 8, 13 and 16 is less well resolved, alongside some of longer internal branches within that subclade. Some lineages, such as the clade currently designated as PeV-3, consists of closely related contemporaneous sequences, whereas other clades, such as the clade designated PeV4, have long internal branching segregating the terminal branching structure.

There are incongruencies in the topologies of the trees reconstructed from the nucleotide and amino acid alignments, including several poorly resolved major lineage placements (Figure 3, Sfigure 4). In the nucleotide tree, PeV-17 clade is a sister clade to PeV-3, with the PeV-10/14 clade in a basal position (aBS 0.82). In the amino acid tree, the PeV-17 clade is basal to the sister clades PeV-3 and PeV-10/14, though this arrangement is less well-supported (aBS 0.55). The PeV-6/18 subtree is basal to the sister clades of PeV-1 and PeV-4/5/8/9/13/16 in the amino acid tree, whereas it is a sister clade to PeV-1 in the nucleotide tree, with both arrangements showing comparable bifurcation support. PeV-13 and 16 are basal to this clade in the nucleotide tree, incongruent with their basal position to PeV-4/5 on the amino acid level, which is marginally better supported (aBS 0.55 vs 0.4). It is clear that additional information is required to accurately resolve the phylogenetic structure of PeVs. However, given the extent of saturation in the nucleotide alignment, it is unlikely to represent a more accurate picture of the evolutionary relatedness of PeVs than the amino acid phylogeny.

To investigate the likely biases introduced by the use of less robust tree reconstruction approaches such as neighbour joining in combination with inadequate evolutionary models or raw distances prevalently used in literature, we used the unedited nucleotide alignment to reconstruct a neighbour joining tree under a Kimura-2 parameter model.^22–28^ This approach severely underestimates the branch lengths in the global phylogeny, as well as resulting in topological incongruencies for multiple lineages relative to the amino acid reconstructed tree (Figure 1B). The inadequacy of phylogenetic reconstruction methods and evolutionary models prevalently used to estimate PeV-A divergence has resulted in several documented incidents of inconsistent typing, including PeV-18 (KT879915) which groups with PeV3 and (KJ796882-3) often annotated as PeV-7, which groups with PeV14.^22^

There is high variability within some of the currently designated genotypes (Figure 1A, 1C, SFigure 5). On both the nucleotide and amino acid level, the more internally divergent genotypes such as PeV-1, 4, and 14 have clear bi- or tri-modal distributions, reflecting long internal branches segregating terminal nodes as observed in the tree. Other distributions e.g. PeV-10 and PeV-15 are skewed by the inclusion of one or two putative genotype members on longer branches.

The genotypes delineated in the current system largely have no spatiotemporal structure, though substantial undersampling limits interpretation. The sampling time frame is not equivalent across genotypes, with sampling globally biased towards the past two decades expectedly (Figure 5, STable 6). The few genotypes (e.g. PeV1 though 5) that span wider intervals have very sparse samples before 2000, with the progenitor strains of PeV-1 and 2 sampled in the 1950s. PeV-A sequences have a very broad and well-mixed geographic distribution, with no clear geographic structure to the currently defined genotypes based on sequence dataset. This is supported by the Wang association index, which showed no significant evidence for geographic structure in the phylogeny of the full dataset (p<0.001). In most countries, multiple genotypes co-circulate contemporaneously, with no clear regional or temporal restrictions to genotypes (STable 5-6, Figure 5, SFigure 6). Globally, PeV-1 and 3 are the predominantly circulating genotypes. Some countries such as Bolivia, India and Ghana, have a disproportionately high number of distinct genotypes circulating relative to the number of available sequences (14, 13 and 12 distinct genotypes respectively, SFigure 7), whereas countries with far higher sampling rates such as Japan, the USA and the Netherlands have five or six genotypes circulating for the equivalent sequence numbers or higher. Australia is the only country with multiple available sequences that all belong to a single genotype, though sampling in Australia is biased by large outbreaks of PeV-3 (STable 5).

**Figure 5.**
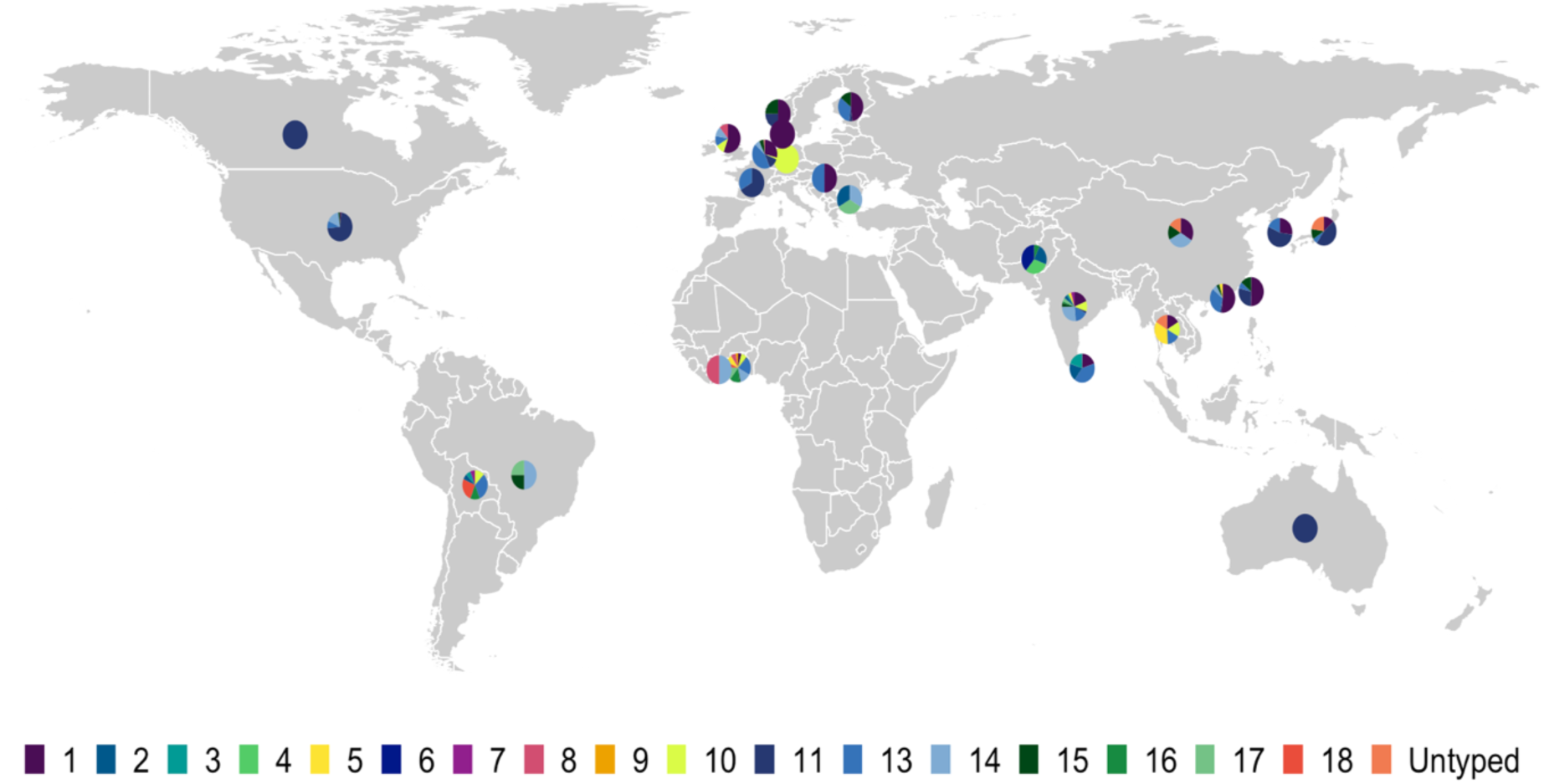
Prevalence of currently designated genotypes by country, indicated by proportion of overall country-level population.

To attempt to resolve possible geographic connections or dissemination networks in the better sampled genotypes, we plotted the geographic origin and date for all highly related virus pairs, defined as pairs of sequences with a pairwise patristic distance below 0.01 substitutions per site in the nucleotide phylogeny (SFigure 8). There appears to be a mild temporal and regional bias in the most closely related sequences, including potential regional networks between Japan, South Korea and Taiwan as well as France and the Netherlands respectively for PeV-3. However, the relatedness of the USA-isolated viruses to the global population supports high levels of geographic mixing, with extreme surveillance biases rendering quantification of these patterns unreliable at this point.

### Evolutionary history of currently defined PeV-A genotypes

The deep divergence between PeV-A lineages, regardless of nomenclature, raises questions around the rate and time scale of the evolutionary history of human parechoviruses, including the divergence dates of the individual lineages.

By root-to-tip regression, there was no evidence of temporal structure across the phylogenies reconstructed from the complete undownsampled and genotype-specific datasets, evident in the extremely low R^2^ values across the different temporal rerooting approaches, where the root is estimated simultaneously with the regression (Figure 6).^66^ Two of the ‘best-fit’ rerooting approaches resulted in negative slopes, interpreted as a negative evolutionary rate. The negative slope was not consistent across different root position optimizations, indicating it is probably a result of the temporal rerooting method mispositioning the root owing to a lack of information. The signal was marginally stronger in the nondownsampled dataset, but still very low. The extent of over-dispersion around the regression line suggests that it may be inappropriate to assume that all branches evolve at the same rate i.e. follow a strict molecular clock model. In several of the rerooting operations, entire lineages e.g. PeV-5 lie below to regression line as clear outliers. We also reduced the sampling timeframe to only include sequences collected after 2006, to investigate the impact of the small number of older anchoring samples on the regression but similarly found a lack of temporal signal for the shorter-term evolution (SFigure 9).

**Figure 6:**
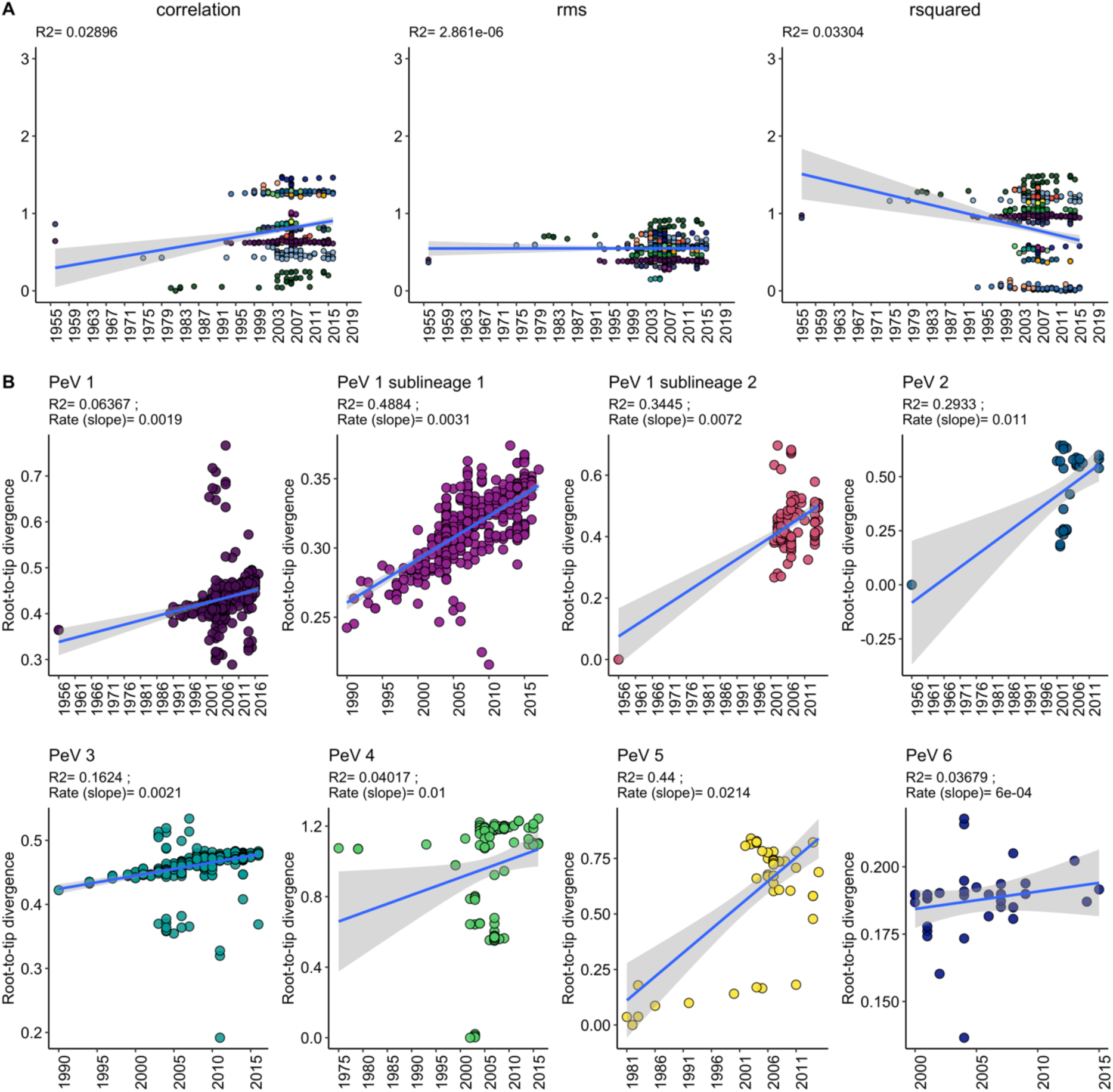
Temporal regressions across re-rooting optimizations. A) Amino acid reconstructed phylogeny of the complete undownsampled dataset. B) Individual PeV-A genotypes as defined by the current genotyping system. RMS = residual mean square.

The high levels of variation around the regression line suggests it would be far more appropriate to assume substantial rate-variation among branches, supported by selection analysis under the aBSREL model, and molecular dating analyses were subsequently conduced with the relaxed uncorrelated log normal clock model. Notably, some PeV-A genotypes including PeV-1 and PeV4-6 have high rates of recombination, whereas negligible rates are reported for PeV-3.^30,67^ Different degrees of recombination among the lineages could result in rate variation. However, using a wide variety of tests provided by the RDP program suite (see Methods), there was no evidence of recombination in the VP1 subgenomic region in the current dataset.

We investigated the temporal structure of the individual currently defined genotypes with sufficient number of samples (n>40), as the currently defined genotypes capture well-defined monophyletic phylogroups. Temporal regression was performed for the undownsampled dataset of PeV-1 and 3, and included the viruses from the test dataset, which were assigned putative genotypes based on their phylogenetic neighborhood.

The temporal signal within the more numerously sequenced genotypes (1-6) was variable (Figure 6). After temporal rerooting to maximize correlation, PeV-1 was structured as two populations with weak temporal signal. The two populations correspond to the two segregated sublineages observed in the phylogeny and have stronger temporal signal when analyzed as two lineages. Genotypes 2, 3 and 5 defined by the current nomenclature system have weak but present temporal signal, with strongly deviating tips in genotypes 3 and 5. Both genotypes 4 and 6 have negligible temporal signal and were not included in the molecular dating analysis. The slope, interpreted as a rough estimation of the rate, was also highly variable across the genotypes. However, root-to-tip regression is inappropriate for statistical hypothesis testing, as the data is correlated owing to shared ancestry, violating the assumption that the data is independently distributed and the current models generally explain little of the observed variance. Outlier tips in genotypes 3 and 5 were excluded from all subsequent analysis.

The substitution rate and tree height (the age of the most recent common ancestor of all samples) estimates for PeV-2 and PeV-5 have very large credibility intervals across models, indicating high levels of uncertainty in these estimates and very slow clock rates (Table 2). This was expected from the temporal regression: the dispersion of the contemporary tips was very high, with the older anchoring sequence inducing the mild linear signal as an artefact, with multiple outliers in PeV-5. The R^2^ signal for PeV-2 weakens to 0.19 when the anchoring sequences from 1956 is removed (data not shown). This emphasizes that signals from temporal regression should be treated with caution, and divergence dating should be subject to prior diagnoses even if runs converge. The rate distribution sampled exclusively from the prior (ignoring the sequence data) also overlaps largely with the recovered rate, suggesting the sequence data may not have sufficient signal to disentangle the tree height and clock rate robustly or recover estimates independent from the specified priors (SFigure 10).^53^ PeV-1 sublineage 2 failed to converged after two combined runs of 500 million steps each.

**Table 2:**
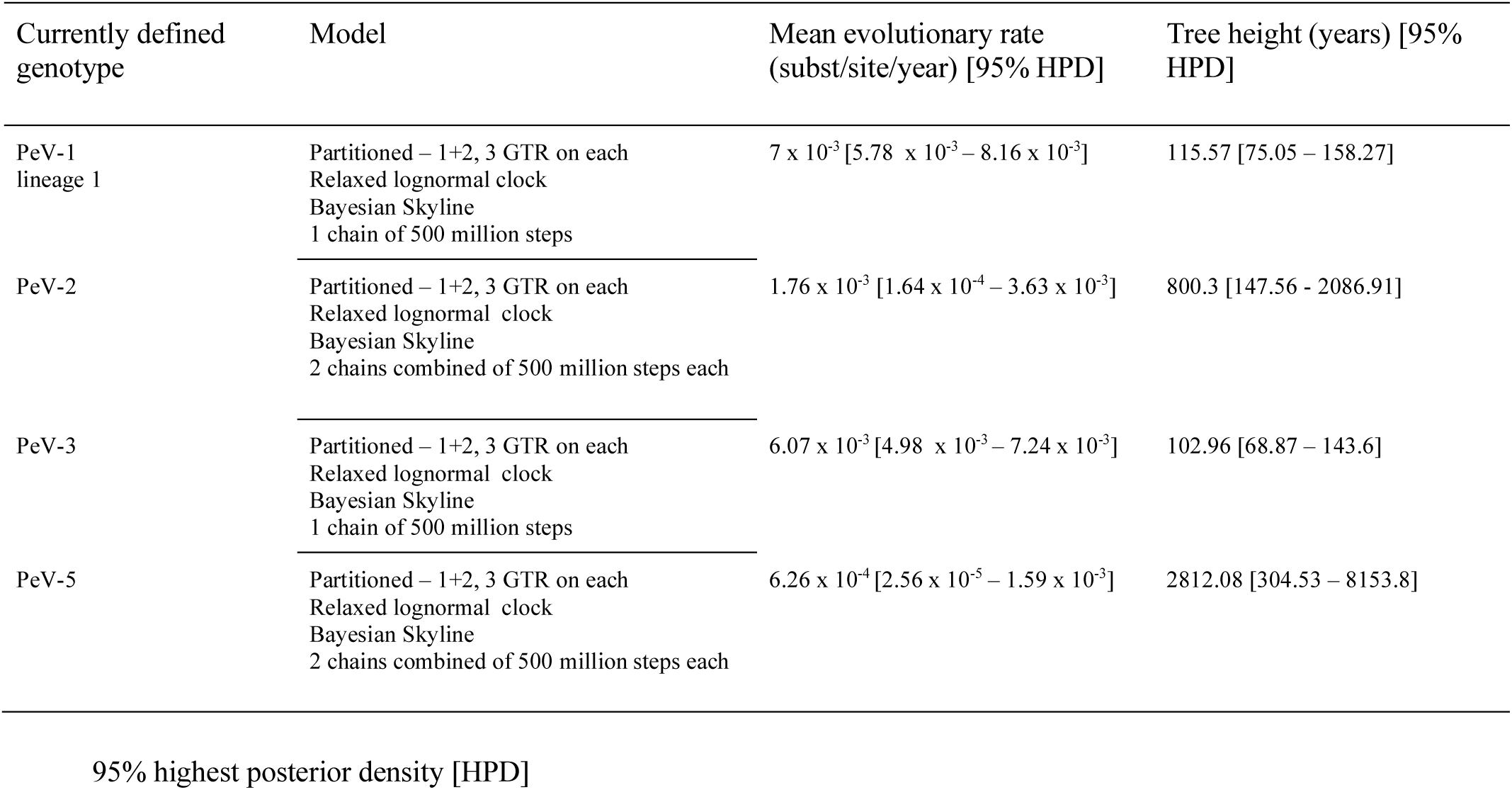
Tree height and evolutionary rate estimates for currently defined PeV-A genotypes

PeV-1 sublineage 1 and PeV-3 have less uncertainty in their estimates. The median tree height, or the age of the most recent common ancestor of all samples, for PeV-1 sublineage 1 and PeV-3 is estimated at 115.57 [95% highest posterior density (HPD), 75.05 −158.27] and 102.96 [68.87 – 143.6] years respectively (Table 2, SFigure 11). The median evolutionary rates for PeV-1 sublineage 1 and PeV-3 fall within the order of magnitude expected of RNA viruses (10^−3^ subst/site/year) but are higher than expected, which may lead to underestimation of the true height of the tree (Table 2).^53,68^

### PhyCLIP-resolved genotyping system

The limitations of the current PeV-A genotyping system and its inconsistent discriminatory information emphasizes the need for a more robust, statistically and phylogenetically informed approach to partition the genetic diversity of PeVs. We used PhyCLIP to delineate evolutionarily relevant genotypes based on phylogenetic relationships of the PeV-A sequences into 26 clusters (Figure 7, STable 7) (compared to 19 in the current system) clustering 97% of all sequences under the optimal clustering configuration (i.e. minimum cluster size of 3, a false discovery rate of 0.2 and a gamma of 3).

**Figure 7:**
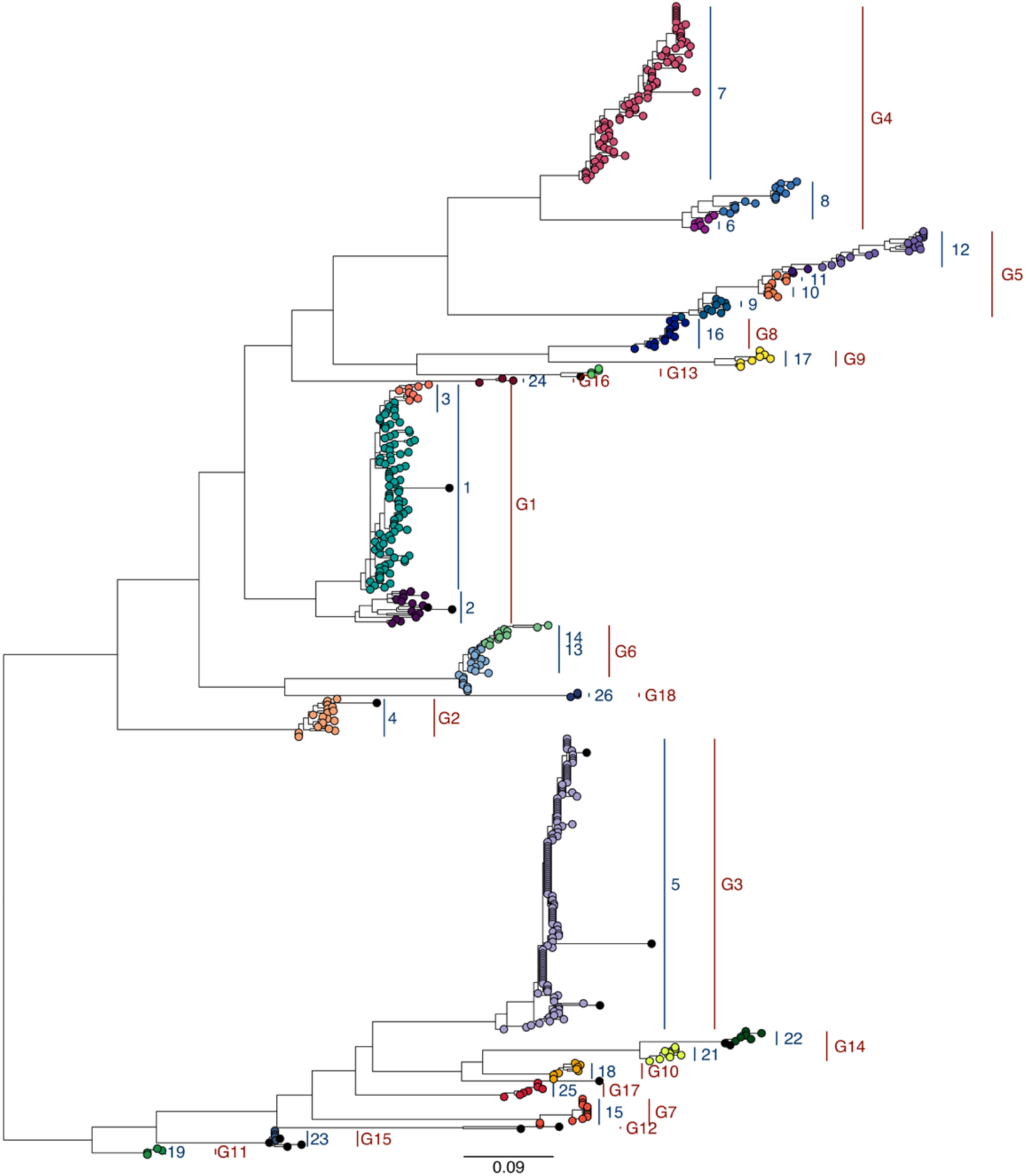
Comparison of PhyCLIP clustering of the phylogeny reconstructed from amino acid alignment of the primary dataset to current nomenclature. Tips are coloured by PhyCLIP designated cluster. Textual annotation indicates PhyCLIP’s clustering in the first set of labels in blue, with the å PeV-A genotype by current system in the second set of labels in red, indicated as e.g. “G1”. Outlier sequences designated by PhyCLIP are indicated in black. See Supplementary table 7 for mapping of current genotypes to PhyCLIP clusters.

PhyCLIP’s cluster topology recapitulates several of the current nomenclature system’s genotypes as single, pure clusters, including PeV-2, 7-10, 11, 13, 15-18 (Figure 7, STable 7). These clusters are sparsely sampled and characterized by low internal divergence, with long interior branches separating them from the rest of the tree. Major genotype PeV-3 is also recapitulated as a single cluster, owing to its distribution of short terminal branches.

Notably, PhyCLIP delineates multiple distinct genotypes in the divergent PeV-A genotypes 1, 4, 5 and 14. For PeV-1, 4 and 14, there are clear phylogenetic separations by long internal branches of the genotypes into two distinct lineages respectively, which PhyCLIP captures (Figures 7, 8). For PeV-1 and 4, PhyCLIP also recovers an additional genotype within each, as the local statistics of the branching pattern supports the initiation of statistically significant divergence at that approximate branch in the lineage. The prototypical PeV-1 Harris strain, which has divergent genetic and antigenic properties from several other PeV-1 strains, forms part of the top PeV-1 lineages.^69^ PeV-5 is phylogenetically structured as one extremely divergent, ladder-like lineage, which PhyCLIP designates into genotypes that step-wise delineate the divergence along the branch owing to its distal dissociation approach.^31^ Even though PhyCLIP increased the clustering resolution within the currently defined genotypes, it did not resolve them into spatiotemporally structured lineages (STable 7) owing to surveillance bias and extensive mixing of PeV-A genotypes globally.

**Figure 8:**
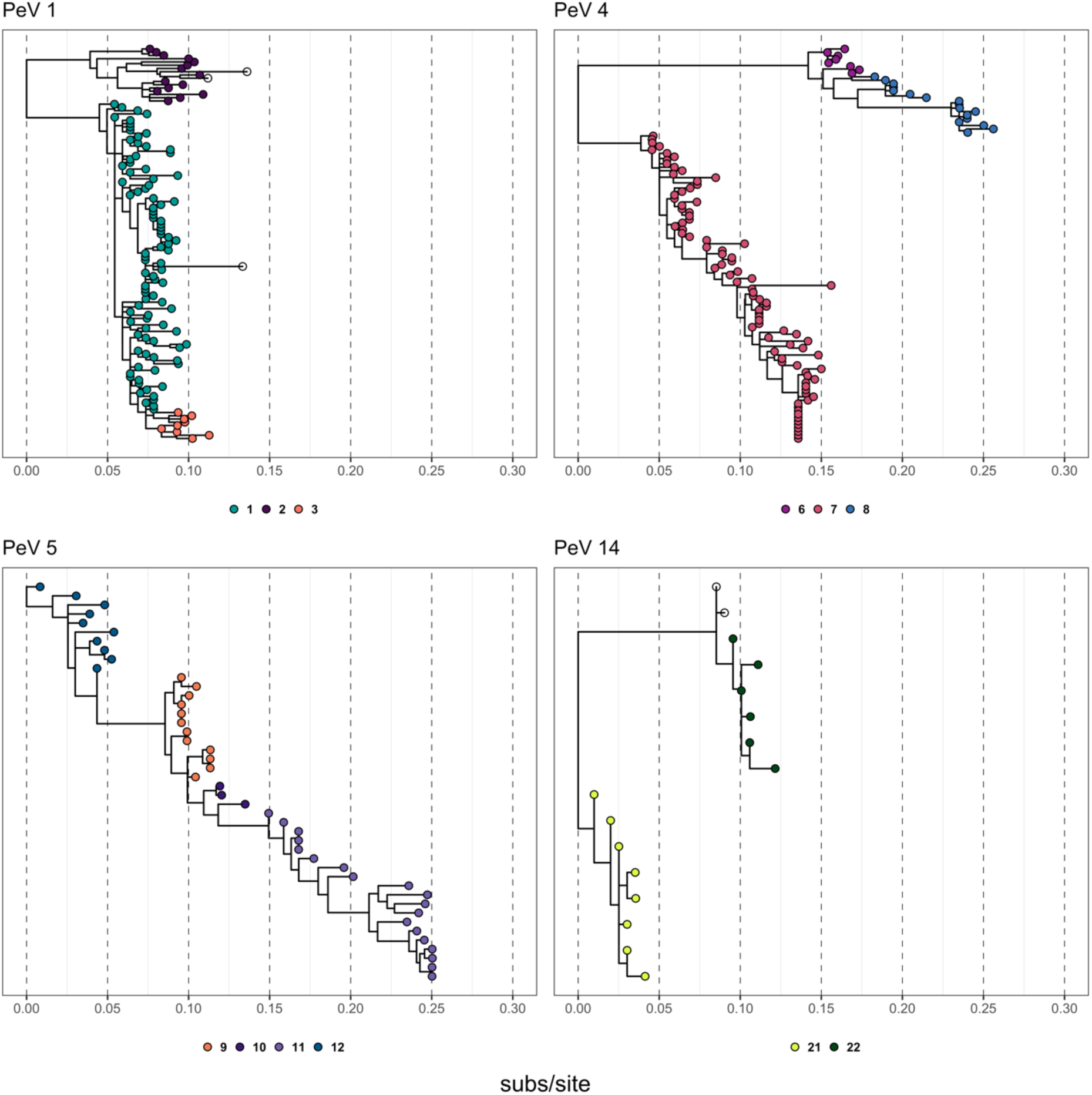
PhyCLIP cluster configuration of the most divergent currently designated PeV-A genotypes 1, 4, 5 and 14. Colored circles below each panel indicate the PhyCLIP genotype.

PeV-12 was designated as unclustered by PhyCLIP, as it is currently only represented by two sequences and fell below the minimum cluster size of 3. PeV-12 was putatively annotated as PhyCLIP cluster 27, to ensure all lineages have a PhyCLIP genotyping identity. Seventeen sequences interspersed through the tree were classified as outliers under PhyCLIP’s distal dissociation approach, which defines an outlier as a sequence that is more than three times the pairwise absolute deviation away from the median patristic distance to the subtending node. Some were classified outliers as a result of the algorithm’s sensitivity in the distal dissociation process, with these sequences showing mild violations of the local branching statistics used to set a divergence limit. This is especially prevalent in regions of the tree with a disproportionately high frequency of identical sequences in small clades e.g. the three sequences in PeV-15. For these sequences, post-hoc absorption into defined clusters is discretionary. Unclustered sequences such as the ones in PeV-1, 3 and 10 show marked divergence to their closest neighbours and are probably true outliers that may represent under-sampled populations or lower quality sequences.

### A new dynamic nomenclature system for PeV-A

The PhyCLIP analyses above provide new genotype identifiers for viruses sequenced up to 2019, but newly sequenced viruses in the coming years will require genotyping. Here, we propose a dynamic nomenclature system that can classify viruses within known diversity as well as be progressively updated to account for both the continual evolution of currently known genotypes and the discovery of new, divergent sequences and lineages. As previously noted, the diversity of PeVs is likely severely underrepresented by sequences in publicly available databases. It is therefore entirely likely that additional surveillance and sampling will reveal extensive undetected diversity, thus shifting the ensemble statistical properties of the inferred phylogeny that inform PhyCLIP’s clustering results. To illustrate this, we subsampled the primary dataset, retaining only viruses collected before 2006 (pre-2006 dataset) and compared its phylogeny as well as its PhyCLIP clustering results to those derived from the primary dataset (see Supplementary Information). Six additional divergent genotypes were detected between 2006 and 2016 (all of which are accounted for in the genotypes shown in Figure 7). Notably, the increased information in the 2016 phylogeny concerning the evolutionary trajectories and diversity of individual genotypes relative to the background diversity progressively improved PhyCLIP’s clustering. Phylogenetic clustering with the PhyCLIP algorithm is therefore a fitting approach to underlie a dynamic nomenclature system.

Optimally, the genotyping tools employed in this nomenclature system would be automated and would not require full phylogenetic reconstruction for every sequence classification, as this will be computationally expensive as the number of PeV-A sequences increases. Phylogenetic placement tools such as RAxML-EPA are ideal for this as they are dependent on a reference phylogeny that incorporates the robust evolutionary models required to more accurately estimate relatedness from sequences as divergent as PeVs while not requiring full phylogenetic reconstruction for every query virus typed.^57^

We broadly categorize the placement results of query viruses into three types with respect to their relative placements against PhyCLIP-defined clusters (Figure 2). Expectedly, untyped viruses that are closely related to viruses within the reference phylogeny will cluster within clusters designated on the reference phylogeny (Figure 2A). These single queries can reliably be genotyped on the reference phylogeny with phylogenetic placement. On the other hand, query sequences that are placed as outliers to designated reference clusters or grouped with outlier lineages designated in the reference phylogeny by PhyCLIP’s distal dissociation would not be nested within known diversity and may warrant the designation of a new genotype (Figure 2B&C). If this type of query begins to dominate with the addition of new sequences, a full phylogenetic reconstruction and clustering should be performed to partition the new standing diversity. Phylogenetic reconstruction is also advisable to reliably resolve the phylogenetic placement of putatively more diverse viruses, as these queries may be placed with lower LWR scores. Full reconstruction and updated clustering will be required if a large amount of sequences is added to databases at once or progressively, as this will shift the global statistics of the underlying dataset PhyCLIP’s operates on, and phylogenetic placement only resolves one query sequence at a time.

Using 123 sequences collected from a recent Malawian cohort study as a test dataset and the primary sequence dataset as reference (see Methods), we investigated 1) if phylogenetic placement could reliably place viruses within known diversity, 2) if phylogenetic placement as outliers to reference clusters or with reference outlier lineages could reliably identify putatively novel divergent lineages or viruses and 3) how the addition of a new, large set of viruses changed PhyCLIP’s reference clustering configuration. The additional sequences from the Malawian cohort were not uniformly distributed across the currently designated or PhyCLIP-derived genotypes (SFigure 12B).

There was a minor topological inconsistency between the phylogeny inclusive of the Malawian sequences (test phylogeny) and the primary phylogeny, with PeV-17/PhyCLIP cluster 25 at an unstable sister clade position to the subtree consisting of PeV-1/PhyCLIP clusters 1-3 in the test phylogeny. However, this bipartition showed very low support (aBS 38). The global patristic distance distribution shifted right significantly relative to the reference phylogeny on the addition of the set of highly divergent viruses from Malawi, increasing the distribution derived within-cluster limit (SFigure 12A). This significant change in the underlying ensemble statistics of the dataset would indicate a nomenclature update is required.

83% of the test sequences were consistently located within a lineage between phylogenetic placement and reconstruction of the test dataset (i.e. query sequences as per the type represented in Figure 2A) with high confidence (average cluster-wise LWR across all queries = 94.9%, standard deviation = 10.8%), indicating accurate phylogenetic placement within known diversity, which is unlikely to require an update (Figure 9, SFigure 13). Divergent sequences are correctly designated as outliers to reference clusters or outlying lineages, which may necessitate phylogenetic reconstruction to confirm placement. The addition of a set of sequences, including divergent sequences, changes the clustering properties with reliable behaviour: divergent sequences are captured as outliers or separate lineages by distal dissociation, which in turn consolidates more closely related clusters (See Supplementary results for details).

**Figure 9:**
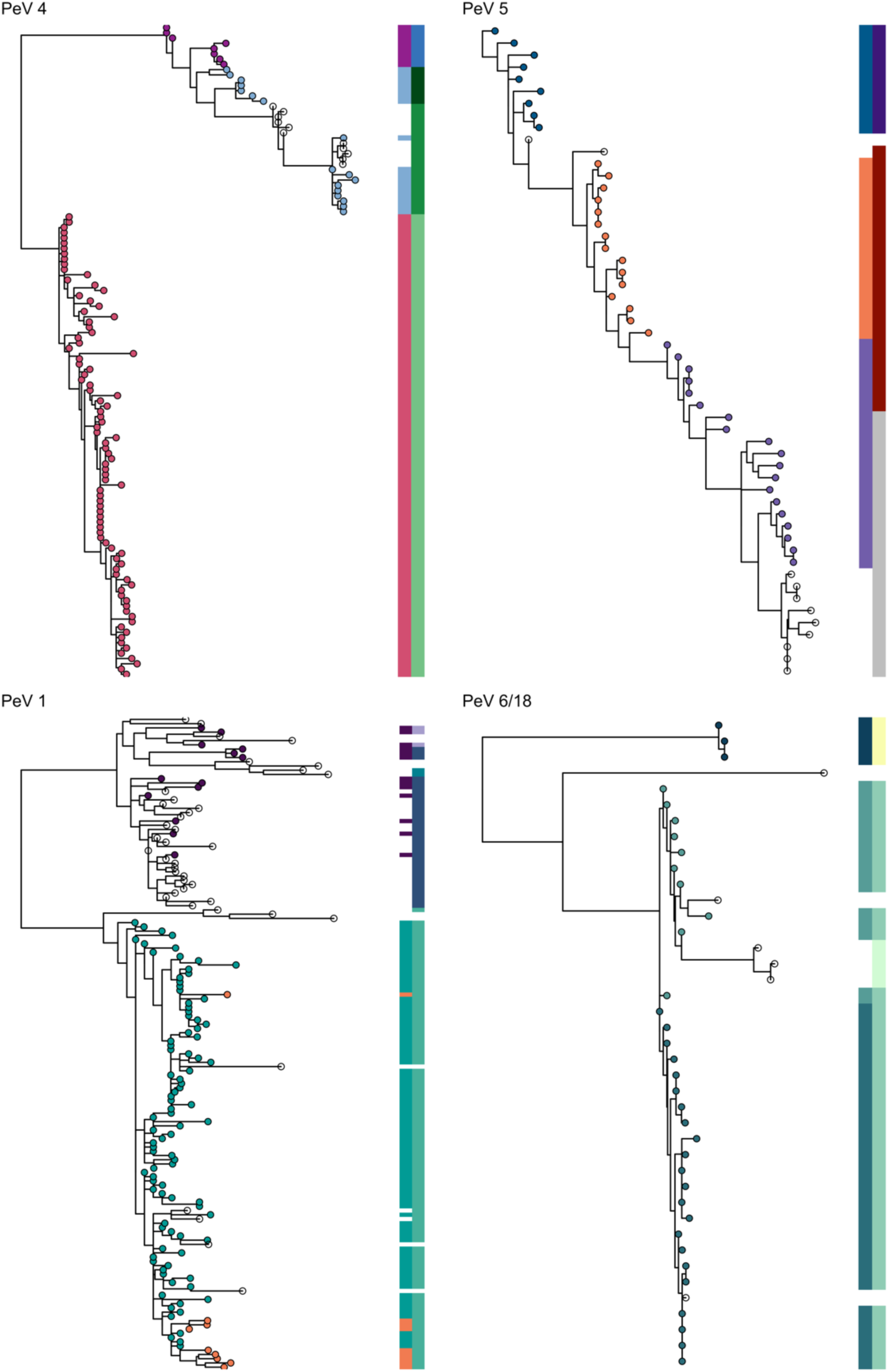
Changes in the clustering topology between optimal phylogenetic clustering of the reference and test phylogenies. Subtrees depicted are from the test phylogeny. Transparent tips indicate the additional viruses. Tips coloured by phylogenetic clustering of primary (reference) phylogeny, which is also the first column of the heat map. Second column is clustering of test phylogeny.

## Discussion

There is a considerable amount of genotypic diversity in PeVs, with notable differences between the currently defined genotypes in terms of their pathogenicity, epidemiology (including the age-distribution of infections), and biological properties, such as receptor usage. However, inference in the genomic epidemiology of PeVs is limited owing to the deep divergence of the lineages and severe substitution saturation in the VP1 gene, which is most commonly sequenced and used for genotyping. There are limited and cautious inferences to be made for the molecular epidemiology of PeVs, each of which is discussed in detail below. 1) Nucleotide-level phylogenetic reconstruction with neighbor joining methods that are currently used for PeV-A genotyping substantially underestimate the evolutionary relationship and the uncertainty thereof between PeV-A lineages. 2) PeVs are substantially undersampled by current surveillance frameworks. 3) It is not possible to recover reliable estimates of the evolutionary history of PeVs with currently available data and tools. Finally, we have introduced our PhyCLIP derived nomenclature system which recovers deep divergences in currently identified PeV-A genotypes in a statistically principled way and provides a reliable basis for genotyping of future PeV-A using phylogenetic placement based on robust, amino-acid level phylogenetic reconstructions. This system recapitulates the currently designated genotypes along long terminal branches, but delineates the internally divergent genotypes more informatively.

### 1. Uncertainty and underestimation in the PeV-A phylogeny require robust phylogenetic reconstruction methods

Our current understanding of the evolutionary relationships between the extant PeV-A lineages is limited by phylogenetic uncertainty in estimation of the deep interior branch lengths and the deep phylogenetic structure. Topological uncertainty owing to artefacts such as long branch attraction introduced by large evolutionary distances restricts our ability to make inferences about the ancestral relatedness of the genotypes. The presence of substitution saturation in the nucleotide alignment emphasizes the need to reconstruct PeV-A phylogenies from amino acid alignments, which are more robust to saturation, as additional caution is required when performing phylogenetic analyses of deep evolutionary time scales in rapidly evolving pathogens, especially on a nucleotide-level.^20,45,58,70^

Additional information is required to resolve the phylogenetic structure of PeVs with more confidence.^71^ Phylogenetic signal from short, subgenomic regions like VP1 is often insufficient, especially as saturation is more pronounced in shorter sequences.^45^ Additionally, the VP1 region is also sometimes just partially sequenced.^23,29,72^ Whole genome sequences could offer valuable additional genetic information to better resolve the phylogenetic relationships of PeV.^73^ However, PeVs have high rates of recombination. With GARD we found 2098 potential breakpoints in the whole genome alignment, and this implies that even with whole genome data unravelling the evolutionary history of PeV-A is likely to be difficult. Increased surveillance and sequencing of quality, full-length VP1 sequences will help resolve the ancestry and relationships between the genotypes.

### 2. PeVs are significantly undersampled by current surveillance strategies

The long internal branches of the phylogeny, both in the deep tree and among more contemporaneous sequences, suggest severe under-sampling of the true diversity of PeVs. Sampling for PeVs is sparse and biased, with differences in study design greatly limiting generalisation and inference. Most available PeV-A sequences are derived from epidemiological studies reporting PeV-A prevalence in cohorts of patients with specific symptoms (e.g. acute gastroenteritis, respiratory or neurological symptoms)^17,22,27,74–79^ and from national enterovirus surveillance programs reporting on PeV-A infections detected in clinical settings.^7,80–82^ The data obtained in these studies and programs are influenced by disease severity, care seeking behavior and the inclusion of specific sample types (e.g. fecal samples, nasopharyngeal swabs or cerebrospinal fluid), and may be biased to genotypes with a higher pathogenicity or a specific tissue tropism. It is thus unclear whether the combination of these passive and active forms of surveillance accurately reflect circulating genotypic diversity.

### 3. Deep evolutionary timescale estimates for PeVs are not possible with available data and tools

Previous molecular clock analysis based on the VP1 region suggests a time to the most recent common ancestor (tMRCA) of 1600.^83^ However, it is likely that these estimates of the evolutionary history of PeVs are substantially underestimated owing to time-dependence in the evolutionary rates, rate heterogeneity among divergent lineages and strong purifying selection along deep-tree branches.^20,84^

The observed genetic diversity of PeVs is likely generated and maintained by a complex interaction of immune dynamics and selection, recombination and co-divergence over deep evolutionary scales. Strong purifying selection acting on most sites results in a high rate of synonymous substitutions as selection functions to constrain changes in the functional sites of VP1.^4–6^ However, there is evidence for diversifying selection in the structured C terminus around the RGD motif and subunit interface region of the VP1 protein as well as suggestively along certain lineages, though statistical power in the current dataset is limited and reduced by conservative multiple testing correction.^1,63–65^ A possible explanation for the deep divergences in the VP1 capsid region phylogeny and the differential presence of the RGD motif could in part be immune-mediated positive selection driving the divergence on a deep evolutionary scale. Use of a different, unknown receptor may result in differences in cell tropisms and the differential diseases severity and epidemiological properties observed for the different genotypes.^85,86^ However, receptor usage for most genotypes has not been characterized.

Estimates of the evolutionary rate and history of PeVs should be treated with caution, as the current full and lineage-specific datasets (excluding current genotypes PeV-1 sublineage 1 and PeV-3) do not have enough information to recover trustworthy parameters robust to prior settings. The extant diversity of current genotype PeV1 sublineage 1 and PeV-3 are estimated to have respective most recent common ancestors within the past two centuries. These two genotypes are also the best sampled, and further sampling may enable more precise molecular dating of the individual additional genotypes, especially if PhyCLIP’s delineation of highly divergent groups proves informative. However, further inference about the evolutionary dynamics of the complete PeV-A phylogeny and the collective divergence dating of the genotypes is limited by the fact that the likely phylogenetic root of PeVs is very deep, with a long evolutionary history of diversification of the genotypes based on the VP1 phylogeny. Resultantly, deep evolutionary relationships among highly divergent viruses may not be resolved with molecular clock analyses that are calibrated on the terminal branches of recent sequences.^20,58,70^

PeVs are RNA viruses, and therefore evolve measurably over shorter timescales owing to substitutions introduced by its error-prone polymerase. It is therefore expected that sequences from a PeV-A dataset serially sampled over 62 years would have accumulated a sufficiently large number of substitution to reliably provide signal for evolutionary history and rate estimation.^66^ However, even if the sampling timescale for PeVs spans 60 years, the sampled time range is highly biased towards the past 20 years and might still be too short relative to the evolutionary timescale of the virus to capture long-term evolutionary processes on those deep time scales.^87^ Time-dependence of rate is particularly prevalent in datasets of deep-time scale pathogens where recent sampling is used to extrapolate back in time. This can lead to bias in molecular dating as shorter, terminal branches have relatively high dN/dS values owing to transient and unfixed substitutions, in contrast to the low dN/dS values for deep interior branches that are under strong purifying selection.^58,84^ Standard GTR+G models do not adequately account for variation in selection pressure across sites and lineages of the phylogeny and will result in under-estimated branch lengths in the presence of strong purifying selection.^47,58,88^ Increased purifying selection at deep evolutionary time scales can lead to an underestimation of the evolutionary history of a phylogeny, as it maintains sequence homology even if synonymous sites are completely saturated.^20,58,84^ Recent work on site-specific models that account for skewed site-specific preferences, both inferred from sequence data and experimentally derived, and the associated saturation have shown marked improvement in phylogenetic fit and branch length estimation accuracy.^89^ However, it is unlikely that there is enough information in the current PeV-A dataset to attempt these parameter-rich models.^90^

The exploratory temporal regression suggests that there is substantial variation in rates (heterotachy) across PeV-A lineages.^70^ This heterogeneity may be due to variation in the strength of selective pressure acting on different lineages, as suggested by the aBSREL analysis. Differential selective constraints may arise from differences in life cycle, including genotype-dependent differential immune pressure and receptor and tissue tropism, resulting in different ratios of synonymous to nonsynonymous substitutions.^89^ Variation of rates may also arise from inherent differences in substitution rates, i.e. differences in synonymous rates, among lineages.^89,91^ The rate variation is unlikely to be a result of different levels of recombination among the lineages, as we did not detect any breakpoints in our VP1 dataset. Rate variation can be modelled with relaxed molecular clock models, which allows for branch-specific rates drawn from an underlying distribution.^25^ However, simulation studies have suggested that a single uncorrelated relaxed clock is not flexible enough to adequately model heterotachy across subtype clades, recovering inaccurate rate estimates and biasing the evolutionary history.^71,91^ Lineage-specific relative rates can also be accommodated with autocorrelated random local clocks, which allows for rate changes on host-specific lineages.^92^ However, these models are biased by definitions of specific lineages and do not account for other sources of rate variation. Recently developed mixed effect clocks combine relaxed and local clocks to model both fixed and random effects in rate variation and have been shown to improve rate estimate accuracy in the presence of heterotachy from mixed sources.^21^ The pronounced sparsity of temporal signal in the primary phylogeny with the presence of purifying selection on a deep evolutionary scale as a confounding factor limited any attempt to reliably estimate evolutionary rates or the dates of divergence in the full PeV-A phylogeny. As estimates of branch length underestimation from the aBSREL model are unlikely to be precise and will have confidence intervals of orders of magnitude, we also refrained from adjusting known estimates of the most recent common ancestor using this information.^20,83^

### 4. New PhyCLIP based genotype nomenclature recovers deep divergences in currently defined genotypes

The current PeV-A nomenclature system falls prey to many of issues described above because of the reliance on neighbour-joining phylogenetic trees and uncorrected genetic distance based thresholds for classifying virus genotypes. The new nomenclature system that we describe here minimises these issues by using the best available phylogenetic methods and the statistically principled PhyCLIP framework to delineate genotypes within the resulting trees. PhyCLIP operates on the long deep and terminal branches of the phylogeny to recapitulate much of the current nomenclature system and to capture clear phylogenetic distinctions in divergent PeV-1, 4, 5 and 14 genotypes. It is unclear if the demarcation of these divergent genotypes into more clusters is informative with regards to the epidemiological or antigenic characterisation of the phylogenetic unit without additional information on serology, life cycle and clinical properties of PeVs. PhyCLIP is sensitive to variation in sampling rates, as its clustering is dependent on the diversity present in the phylogeny. PhyCLIP will perform optimally when the background diversity of the population tested against is comprehensive or well representative. The evolutionary continuum of genotypes PeV-1 and 3 is far better approximated in the current dataset than less sampled genotypes. Nonetheless, the long interior branches separating deep and more terminal clades enables PhyCLIP to delineate clusters.

The dynamic nomenclature system combining progressive clustering by PhyCLIP with phylogenetic placement shows promise. Phylogenetic placement with RAxML-EPA accurately places sequences with close relatives within existing diversity and is capable of correctly designating divergent sequences as outliers to reference clusters. The addition of a number of divergent sequences or a large set of sequences to the existing diversity will likely require that our PhyCLIP based pipeline be re-run as PhyCLIP specifically operates on the underlying distribution of genetic divergence in the phylogenetic tree. The exact timing of such a re-evaluation will depend on the rate of new sequencing data generation.

## Acknowledgements

The authors would like to thank Sergei L. Kosakovsky Pond and Joel Wertheim for their help.

