## Supplementary information for "Genotypic diversity and dynamic nomenclature of *Parechovirus A*"

#### Supplementary Methods

##### Wang association index for phylogenetic structure of traits

The Wang association index measures whether a trait is more phylogenetically structured than expected by chance, suggesting possible correlation of traits by shared ancestry.[54,55] The association index quantifies the strength of association and is calculated as per equation 1:

$$\mathrm{AI} = \sum_{i=1}^{k} \frac{1-f_i}{2^{m_i-1}} \tag{1}$$

Where $f_i$ *is* the frequency of the most common trait (here geographic region or country) subtended by node *i* and $m_i$ is the number of tips node *i* subtends. Summarized over all internal nodes, a low AI represents a strong phylogeny-trait association. P-values are obtained from 1000 permutation tests, where the AI is calculated after traits are randomized across all tips in the tree. Phylogenetic error is usually incorporated by average the AI across a posterior distribution of phylogenetic trees, or with e.g. 100 maximum-likelihood trees reconstructed from bootstrapped alignments. However, the phylogenetic requirements for reliable phylogenetic reconstruction from the primary dataset and the associated computational burden precluded this. The Wang association index and permutation testing were implemented with a custom python script.

#### Supplementary Results

##### Sensitivity of PhyCLIP phylogenetic clustering to sampling

The long interior branches of the phylogeny suggest severe under-sampling of the true diversity of HPeV (SFigure 4). Additional sampling of divergent HPeV lineages will most likely break down or affect these long internal branches, which might help resolve the phylogenetic structure. Concurrently, a more complete picture of the diversity of the HPeV population will fundamentally shift definitions of what are considered sufficiently closely related viruses. The dynamic is central to PhyCLIP, as the addition of a large number of potentially divergent viruses to the underlying data will shift the ensemble statistical properties of the phylogeny, driving changes in the clustering configuration as the global patristic distance distribution, topology, and statistical power between data sets is altered. To investigate this sensitivity to sampling, we downsampled the primary phylogeny to only include viruses sampled before 2006 and compared the overall phylogenetic inference as well as PhyCLIP clustering configuration.

From 2006 onwards, 420 sequences were added to the primary tree. This included the addition of all of the sequences representing the divergent lineages demarcated as genotypes 8, 13 and 15-19. This resulted in incongruencies in the deep topology between the primary and pre-2006 phylogenies, which may drive changes in clustering configurations between phylogenies. The two major clades are maintained, with no instabilities in the ancestral trace of the clade encompassing HPeV 3, 7, 10, 11, 12 and 14. The additional of HPeV 18 and notably HPeV 16 changes the branching order of the other major clade. Additionally, HPeV 6 is placed as sister clade to HPeV 1 in pre-2006 tree (aBS support of 97), whereas it sits basal to it in the full phylogeny (aBS support 78). HPeV 2 is placed basal to the sister clade HPeV 1 and 6 in the pre-2006 phylogeny (aBS support 89), which is incongruent with its placement as basal to the major clade in the full phylogeny (aBS support 100).

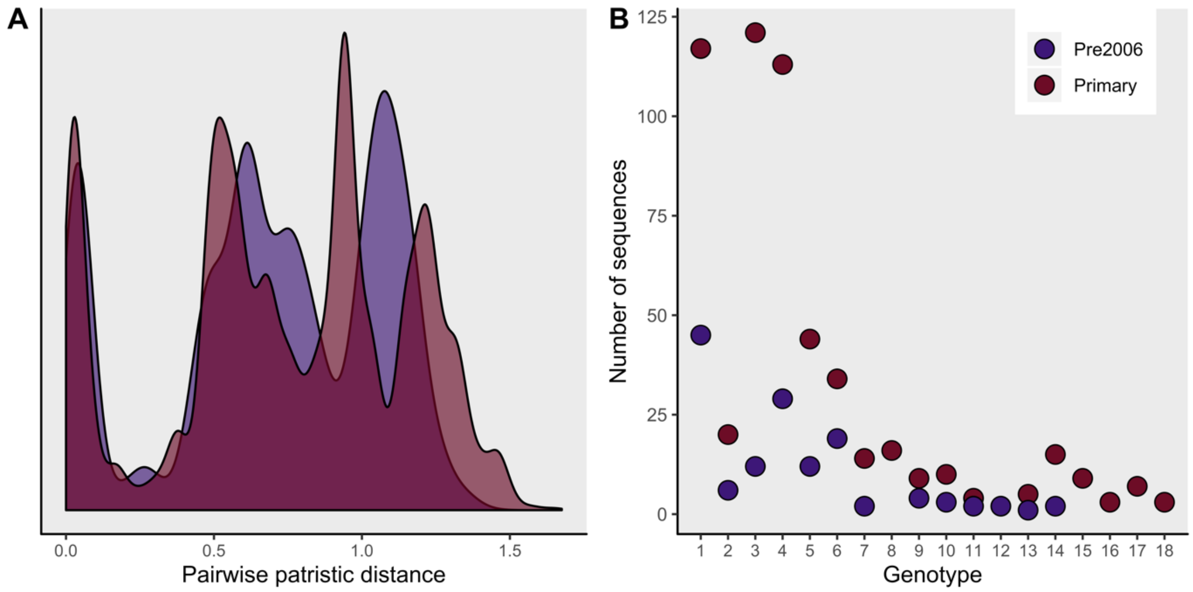


Figure 1.1: A) Density of pairwise patristic distance distribution of the primary and pre2006 phylogenies B) Genotype distribution of the number of sequences by genotype.

The addition of a set of highly divergent sequences in currently defined genotypes 8, 13 and 15-19 increased the spread of the global pairwise patristic distance distribution from the pre2006 to primary phylogeny (Figure 1.1). However, the majority of additionally sampled viruses were from lower internally divergent genotypes such as HPeV 1 (lineage 1) and 3. This caused a relative shift to the left in some peaks, concurrently biasing the within-cluster limit derived from the distribution downwards relative to the pre-2006 within-cluster limit. This would expectedly decrease tolerance of allowable within-cluster divergence.

The clustering configurations of genotypes 2-4, 7, 10 and 11 were stable between the two phylogenies (Figure 1.2). This is expected, as these genotypes have low internal divergence, and are separated from the rest of the phylogeny on long interior branches. In the pre-2006 tree, HPeV 1 was not clustered into the two divergent lineages resolved in the full phylogeny. The smaller lineage was undersampled relative the larger lineage in the pre-2006 phylogeny and relative to its representation in the full tree. As the more divergent HPeV 1 lineage 2 sequences were not present in the pre-2006 phylogeny, the two lineages were consolidated as the second lineage was not yet distinct enough from the larger lineage 1. HPeV 6 is also captured as a single cluster pre-2006, as the diversification branching pattern up top is undersampled prior to 2006. The internally divergent HPeV 5 and 14 were severely under-sampled before 2006, resulting in the minor of the two divergent lineages resolved in the primary phylogeny for each genotype being left unclustered, as they statistically present as outlier lineages. Sequences from genotypes 12 and 13 are also designated outliers, as the intermediate branches are undersampled in the pre-2006 tree.

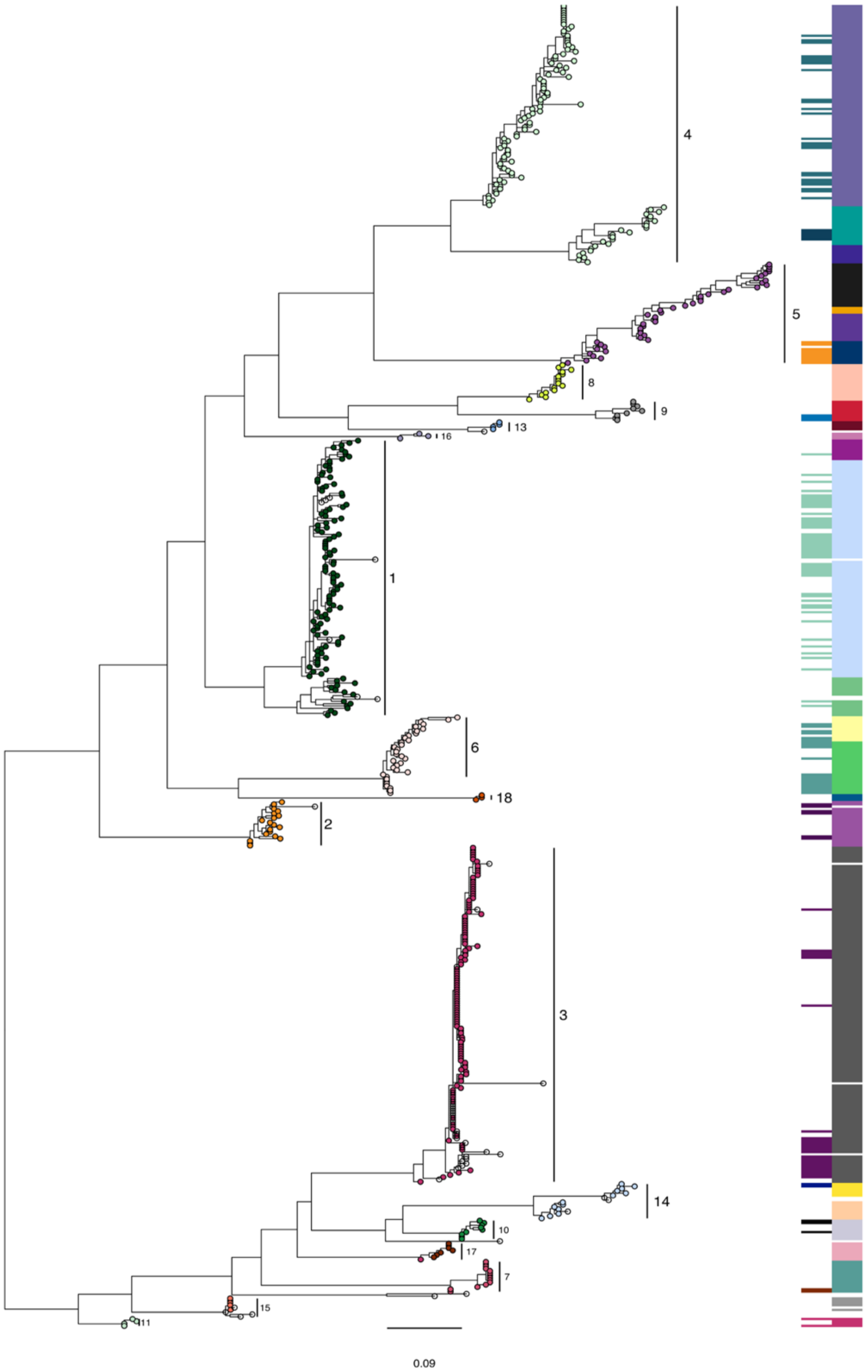


Figure 1.2: Clustering comparison of PhyCLIP results on the pre2006 (column one of the heatmap) and primary phylogenies (column two). Tree tips are coloured and annotated by current genotyping system.

**Accuracy of phylogenetic placement and influence of placed sequences on PhyCLIP's clustering properties**

Six viruses were placed as outliers to the PhyCLIP reference cluster 2 (aka cluster 2 defined in the primary phylogeny) by phylogenetic placement. In the reconstructed test phylogeny clustered by PhyCLIP, the inclusion of these more divergent viruses as well as the additional Malawian sequences correctly placed within the known diversity of PhyCLIP cluster 2 shifted the boundaries of the local statistics to resolve four clusters (Figure 11). One cluster was formed almost exclusively from the test set viruses, and encompassed four of the phylogenetic placement outliers, while the other two outliers formed their own cluster. Two phylogenetic placement outlier sequences were placed on the branch separating the unclustered HPEV-2 progenitor from the rest of HPEV-2/ PhyCLIP cluster 4 in the reference clustering. Both sequences were consolidated into the cluster in the updated phylogenetic clustering of the test phylogeny, while the progenitor remained unclustered. Four sequences were placed on one of the branches in HPEV-12/PhyCLIP cluster 27 of the reference phylogeny. These viruses were congruently clustered as a divergent descendant cluster of the reference HPEV-12/PhyCLIP cluster 27 sequences in the updated test phylogenetic clustering. Test set sequences that were placed within the known diversity of reference clusters representing HPEV-6/PhyCLIP cluster 14, HPeV14/PhyCLIP clusters 21 and 22 and HPeV1/PhyCLIP cluster 1 respectively were designated as outliers in the updated phylogenetic clustering of the test dataset. In all cases, the sequences were basal to the cluster it was purportedly part of and separated by a long branch. Topological inconsistencies between full reconstruction and phylogenetic placement may be ascribed to differences in the models RAxML-EPA implements relative to the more robust reconstruction under comprehensive models in IQtree.

There were other expected changes in the clustering topology between the reference and test phylogeny, driven by the significant changes in the underlying dataset (Figure 11, SFigure 11 ). For HPEV-4, some of the Malawian viruses are placed on the long interior branch separating a previously delineated but divergent reference cluster (PhyCLIP cluster) 8. The additional information resolves the cluster into two in the test phylogeny. The addition of a clade of viruses to the terminus of the very divergent HPEV-5 lineage, representing PhyCLIP clusters 9-12, resulted in the consolidation of three clusters in the reference phylogeny into two in the test phylogeny, with a boundary shift for a few viruses in the intermediate branches as the additional diversity makes them appear less internally divergent. The addition of a nested divergent lineage to HPEV-6/PhyCLIP 13 resolved an additional lineage, consolidating the previous two clusters into one. In the full phylogeny, the local statistics showed statistically supported divergence branching in the top lineage (PhyCLIP cluster 14), but this fails testing when the additional lineage is incorporated into the local statistics. One sequence (MH339678) isolated from the Malawian cohort designated as a putative genotype 19 strain (LWR=75.5%) is separated from HPEV-6/PhyCLIP cluster 13 on a very long branch (0.65 expected substitutions per site), supporting its designation.

### Supplementary Figures

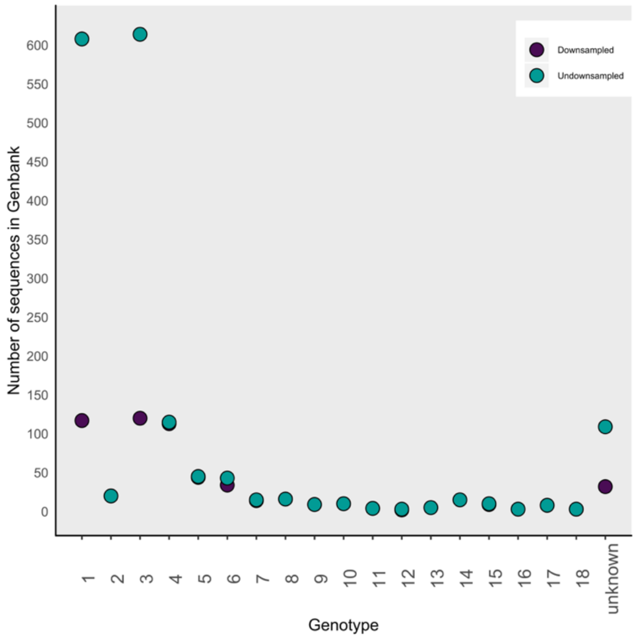


Supplementary figure 1: Distribution of publicly available sequences by genotype on Genbank. Genotype assigned based on Genbank metadata or annotation in associated literature.

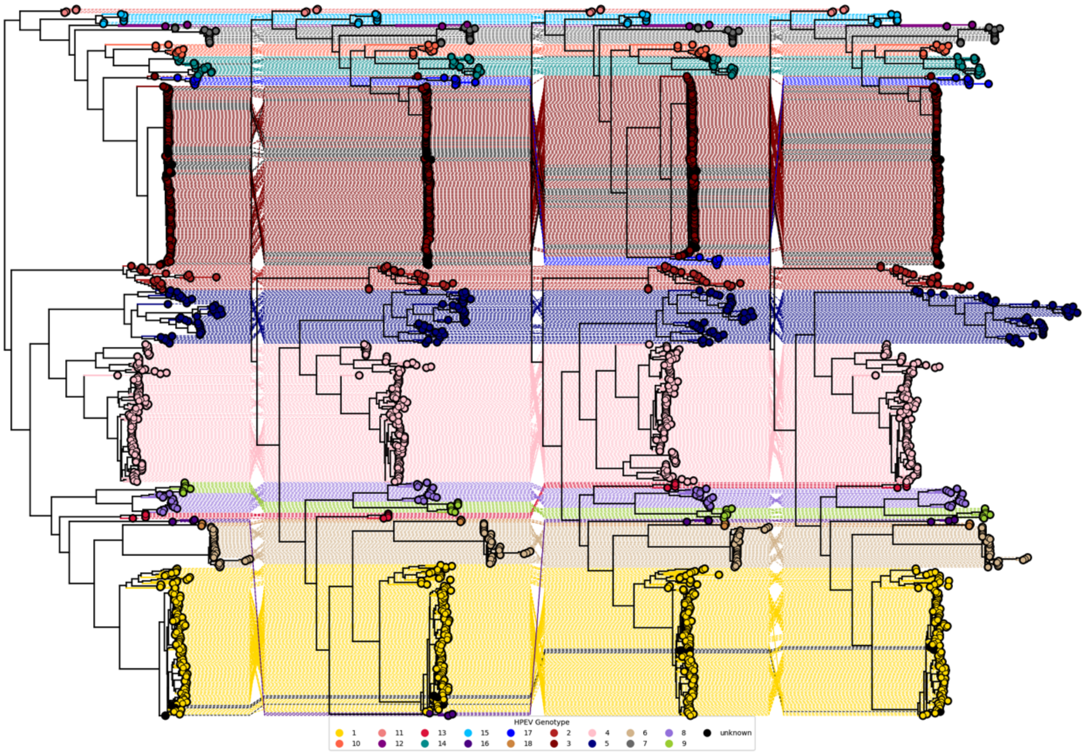

Supplementary figure 2: Phylogenetic incongruencies across phylogenies reconstructed from the different nucleotide alignments. The phylogenetic trees were reconstructed from the alignments in order: unedited, GBlocks, trimmed, Trimal (See Methods). Apart from inter-genotype incongruencies and nodal rotations, the only major discrepancy is the placement of genotype 13.

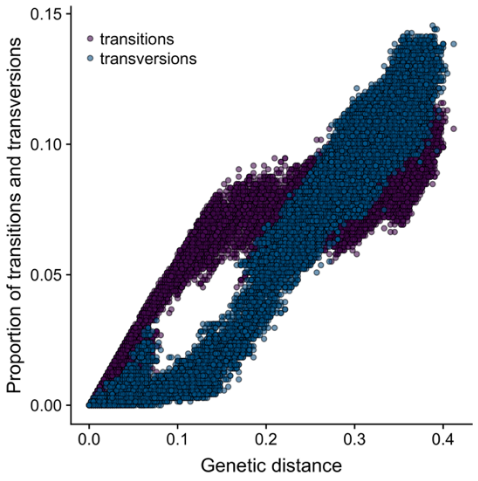


Supplementary figure 3: Frequency of transitions and transversion plotted against genetic distance for the unedited nucleotide alignment. To investigate the extent of saturation in our datasets, we plotted the transition and transversion frequencies against genetic distance (Sfigure 3). If there are high levels of saturation underlying the sequences, the transversion frequencies should be higher than the transition frequencies for larger genetic distances, as observed.[62]

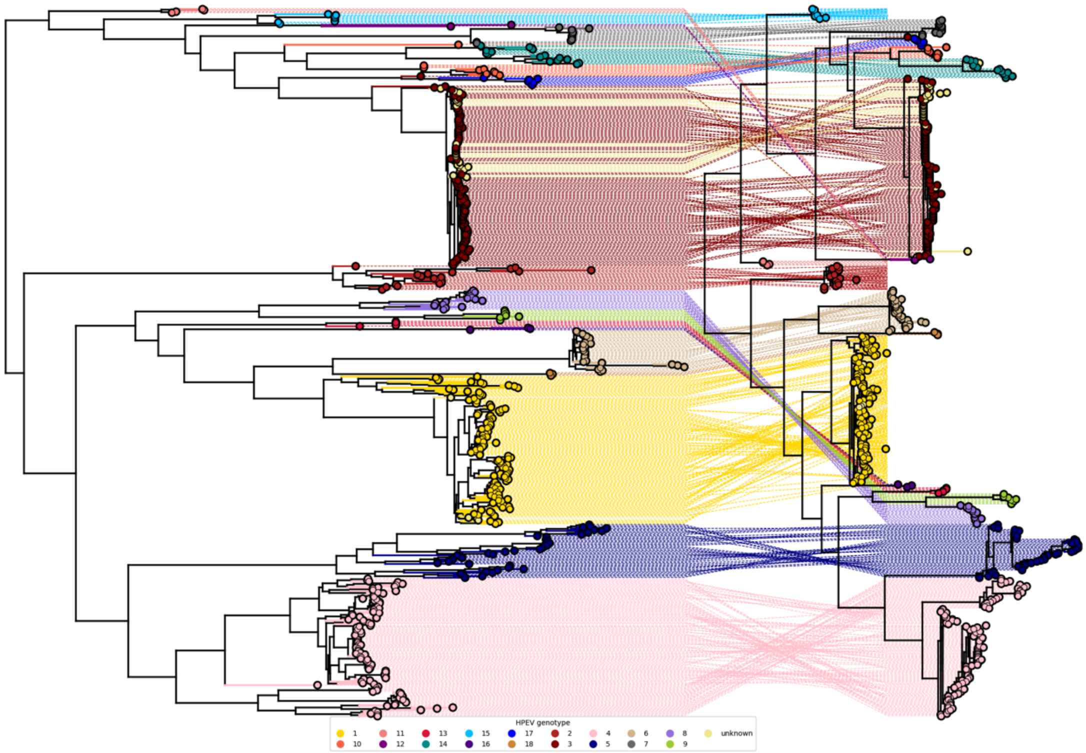


Supplementary figure 4: Incongruencies between phylogenies reconstructed from nucleotide and amino acid alignments

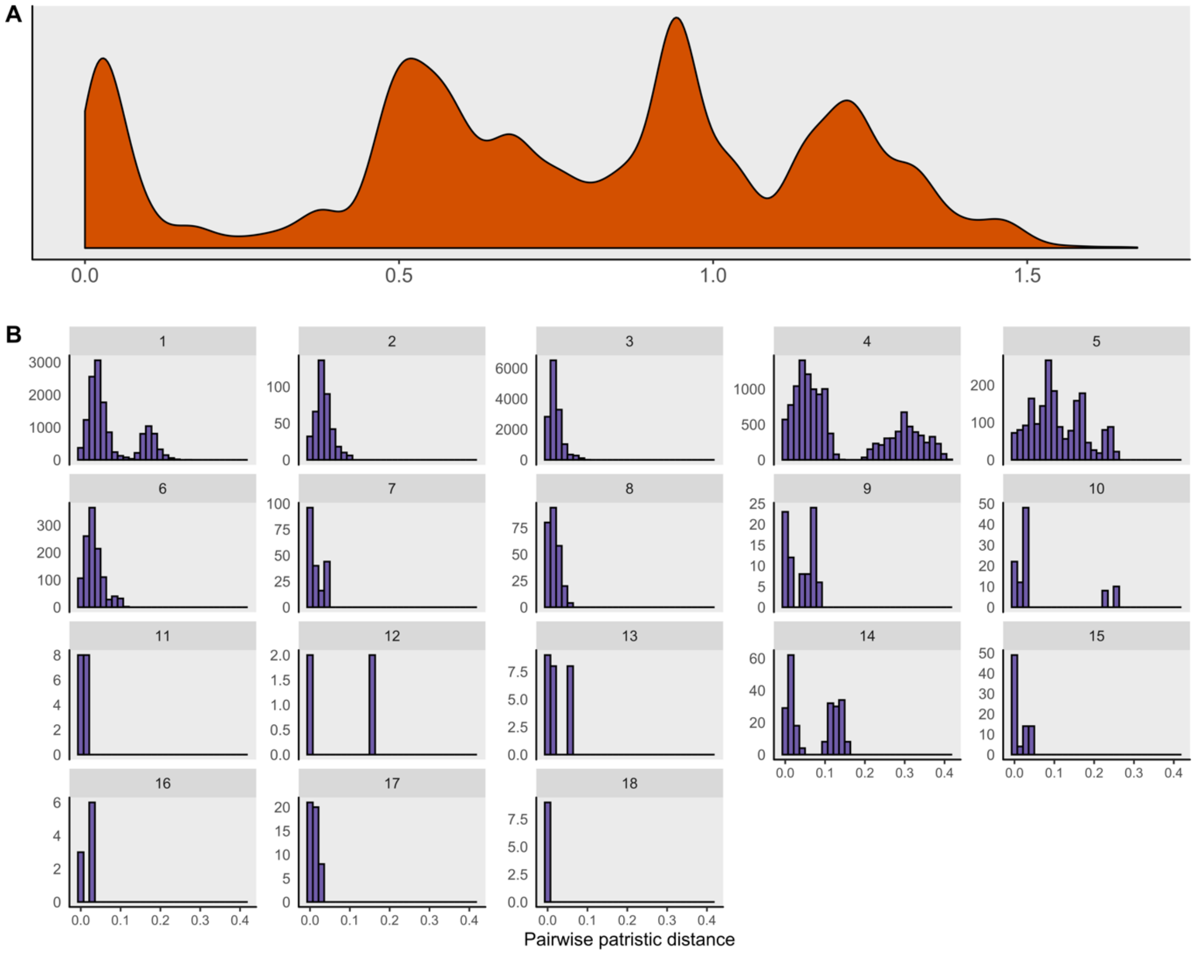


Supplementary figure 5: A) Density of pairwise patristic distance distribution in the reconstructed amino acid phylogeny. B) Within-genotype pairwise patristic distance, as designated by the current nomenclature in the primary amino acid phylogeny

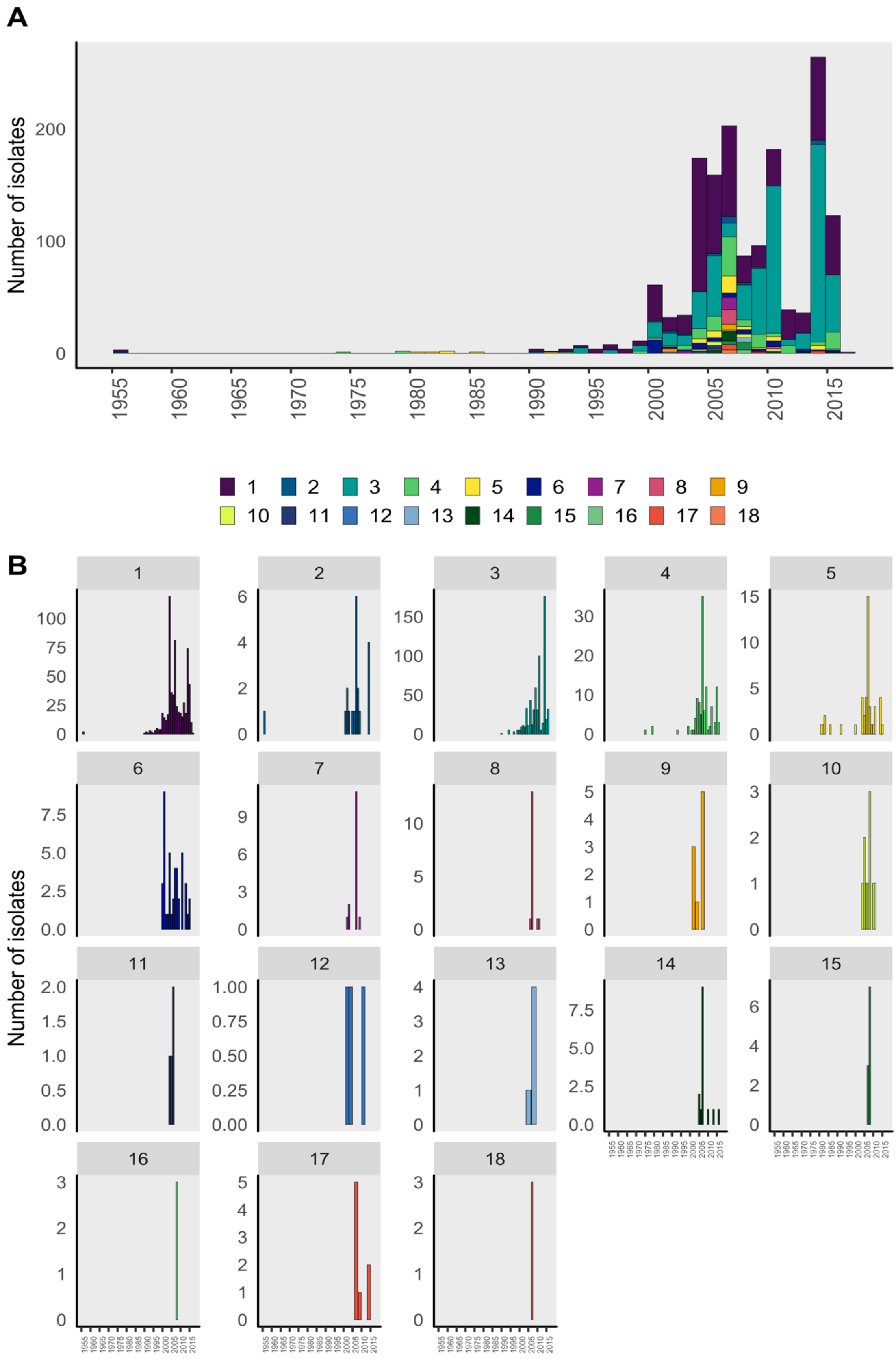


Supplementary figure 6: Distribution of sampling times of viruses in Genbank by designated genotype.

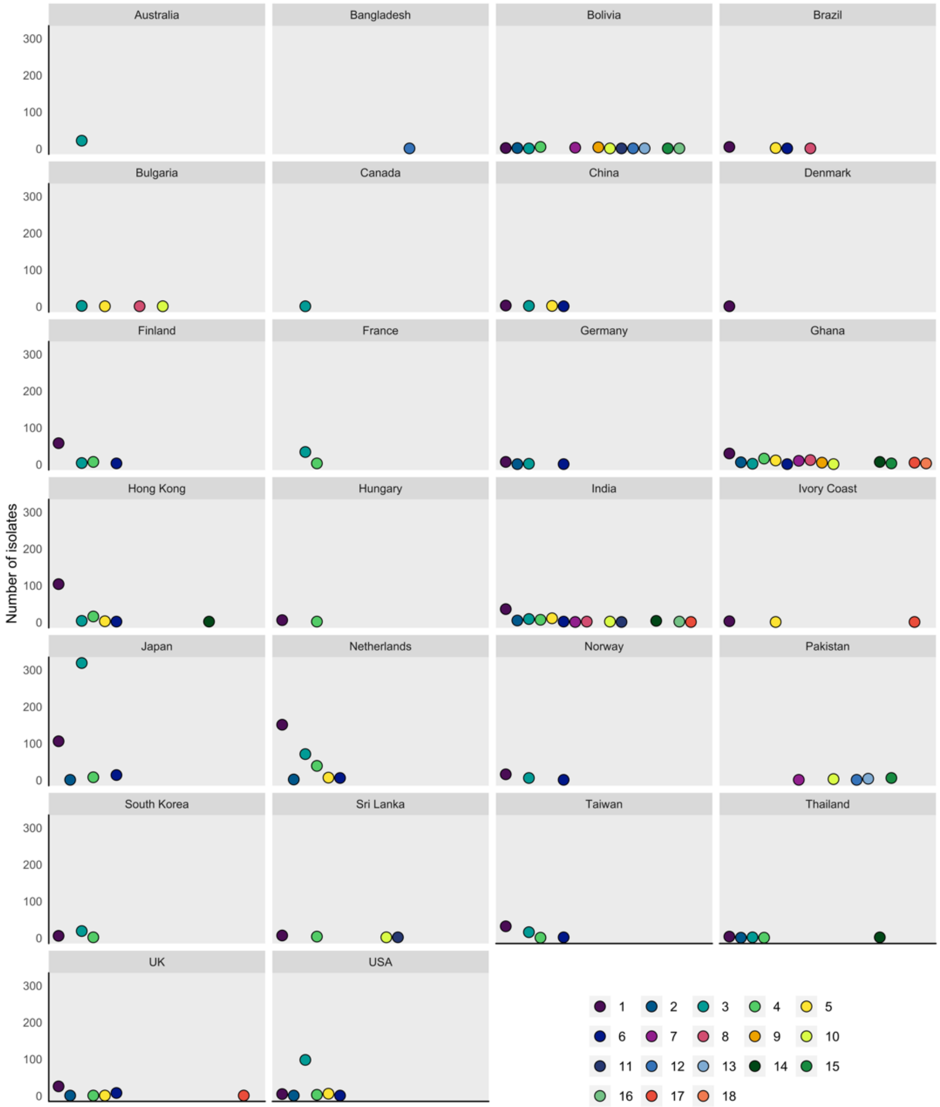


Supplementary figure 7: Distribution of genotypic diversity across countries

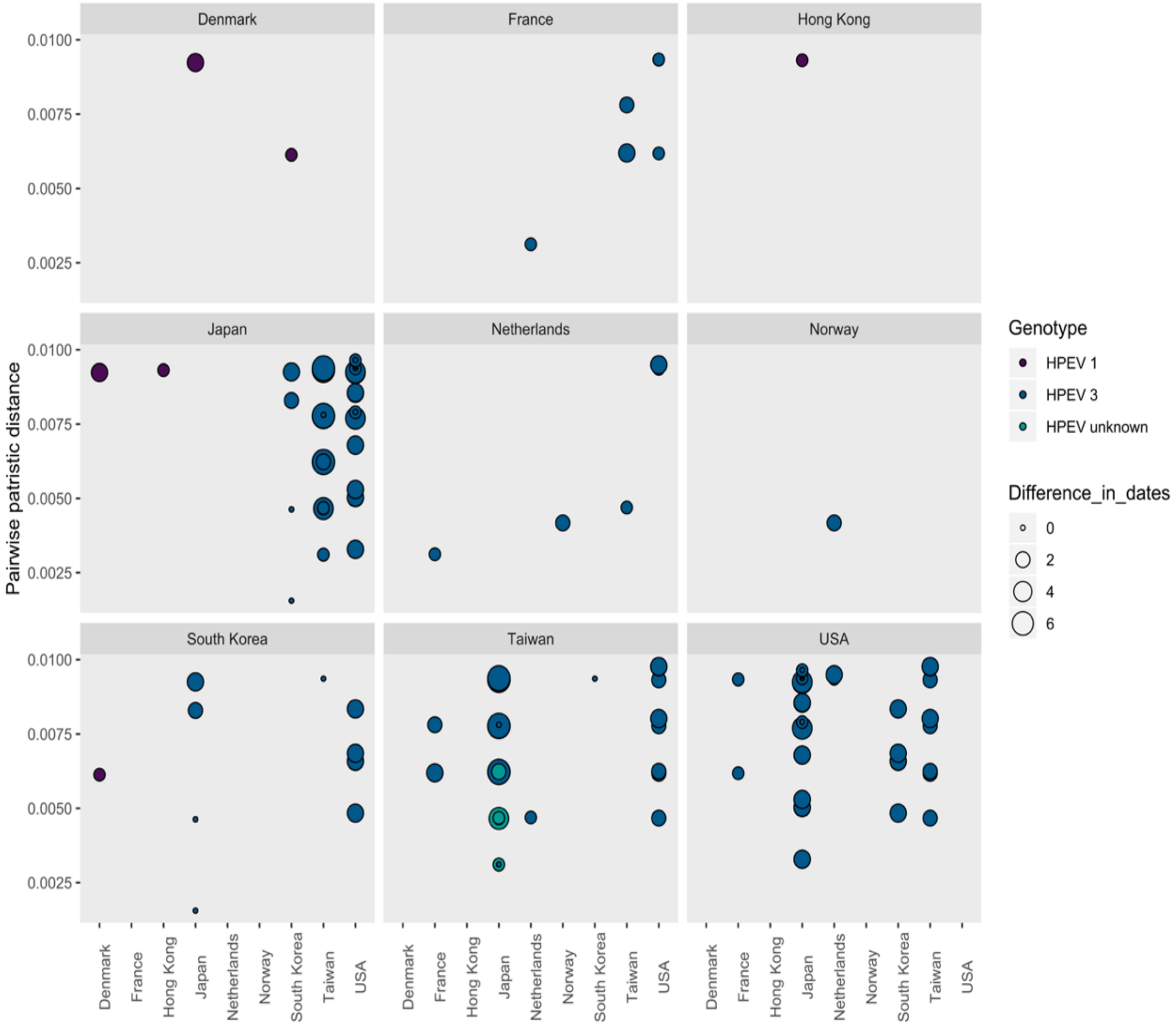


Supplementary figure 8: Migratory networks of closely related sequence pairs (i.e. their pairwise nucleotide patristic distance <0.01 substitutions per site). Size indicates difference in sampling dates in years. These distances are unlikely to be largely inaccurate as short branches were fairly accurate in the aBSREL re-estimation (Figure 4).

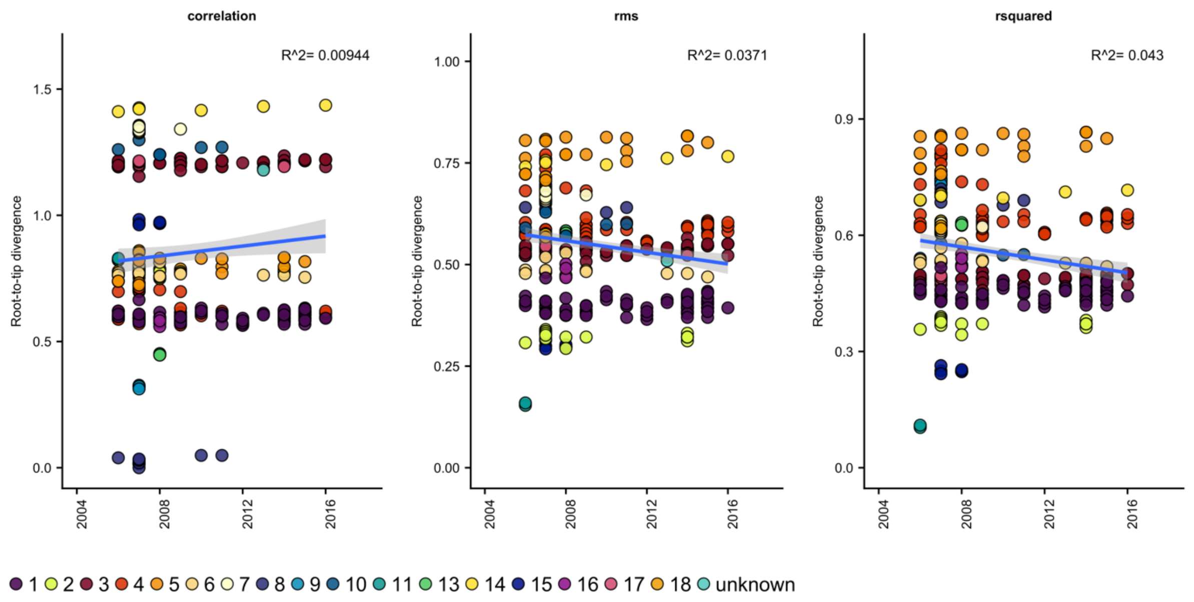


Supplementary figure 9: Temporal regression of phylogeny constructed from all sequences after 2006, rerooted with root optimization approaches. Colours indicate HPeV genotype. RMS = residual mean square.

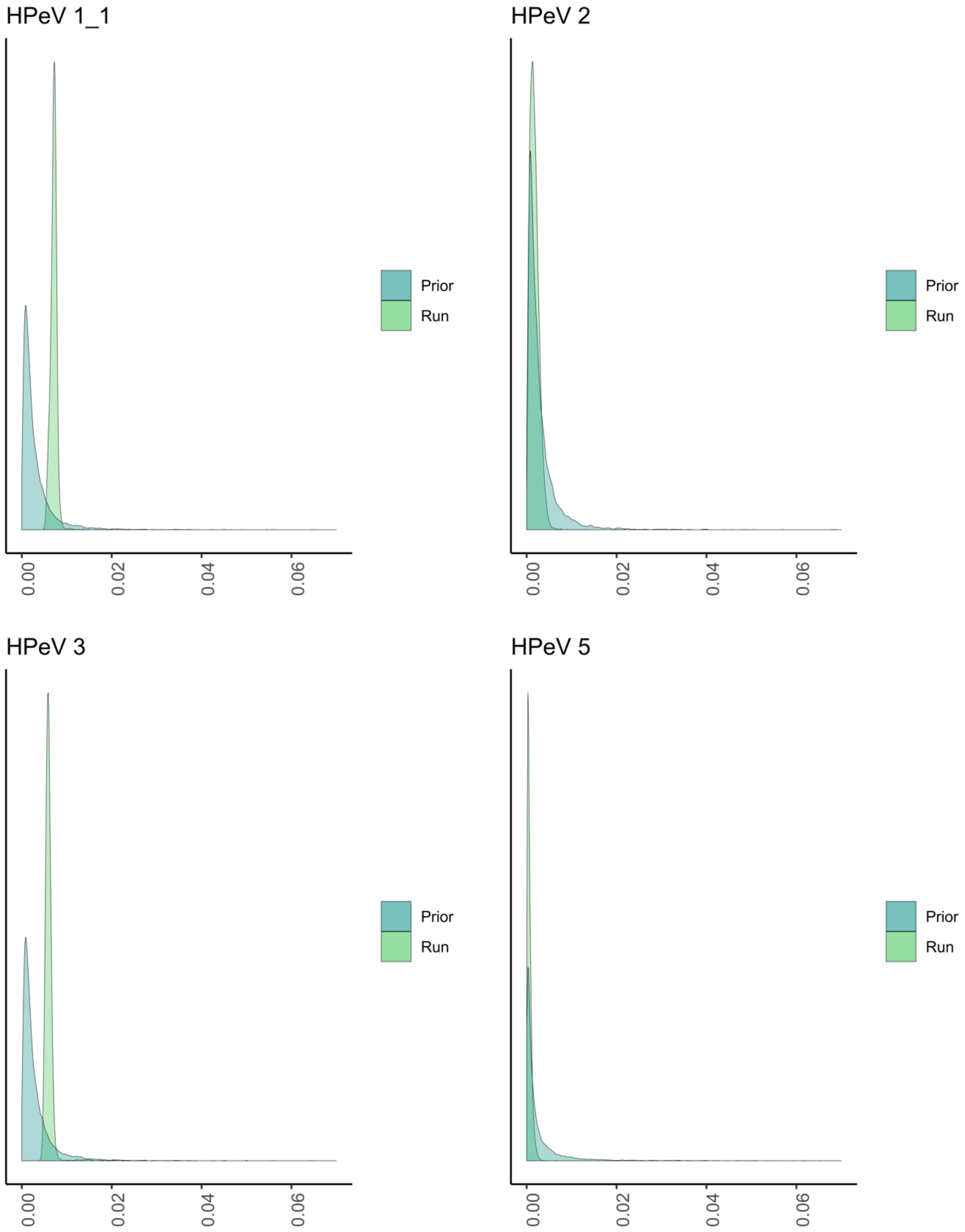


Supplementary Figure 10: Prior diagnoses of the clock rate distribution of HPeV genotypes for the model with the Bayesian skyline tree prior. X-axis was truncated to 0.07 for clarity.

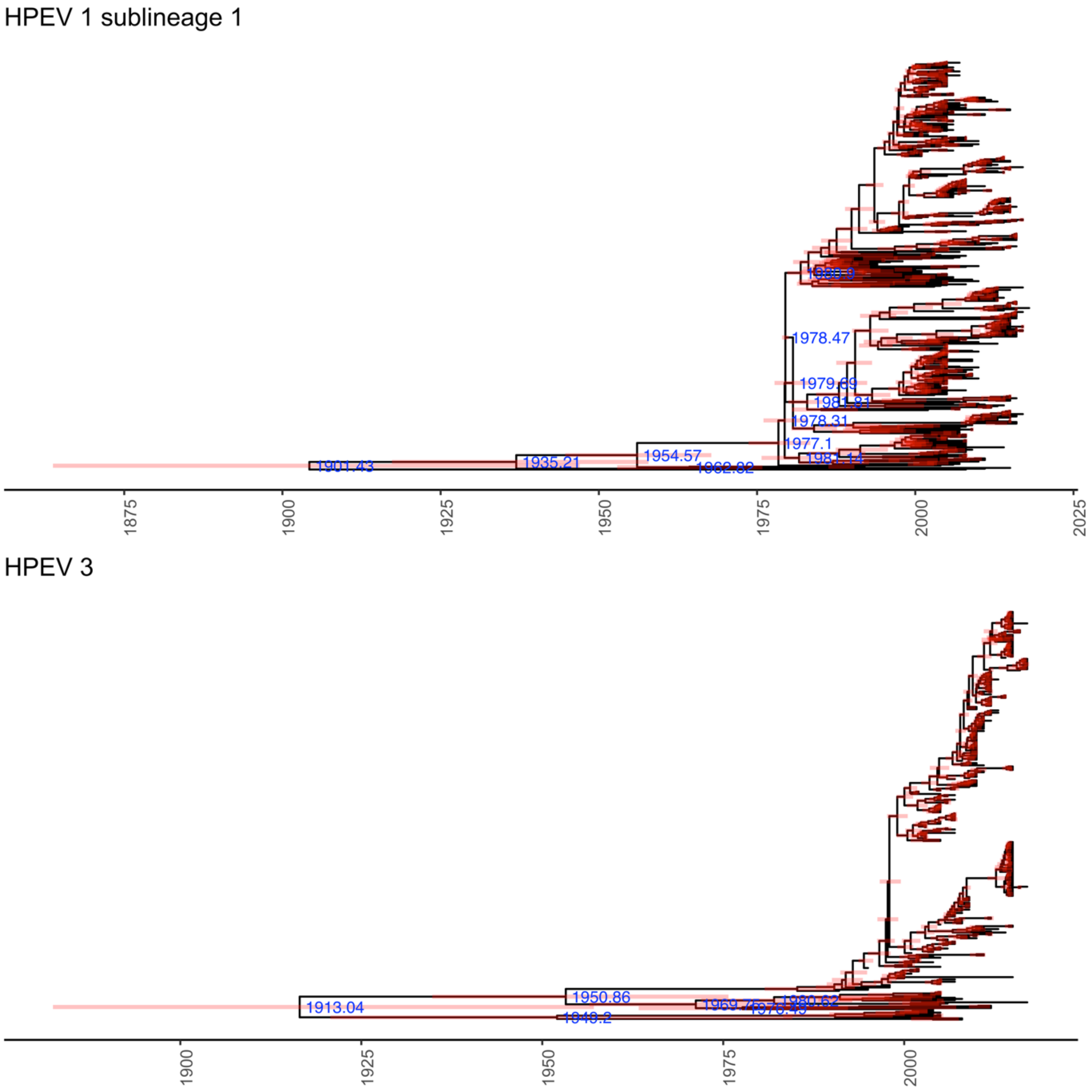


Supplementary Figure 11: Time-calibrated phylogenies of currently defined HPeV 1 sublineage 1 and HPEV 3. Red bars indicate 95% HPD interval around node age, with all nodes older than 35 years annotated.

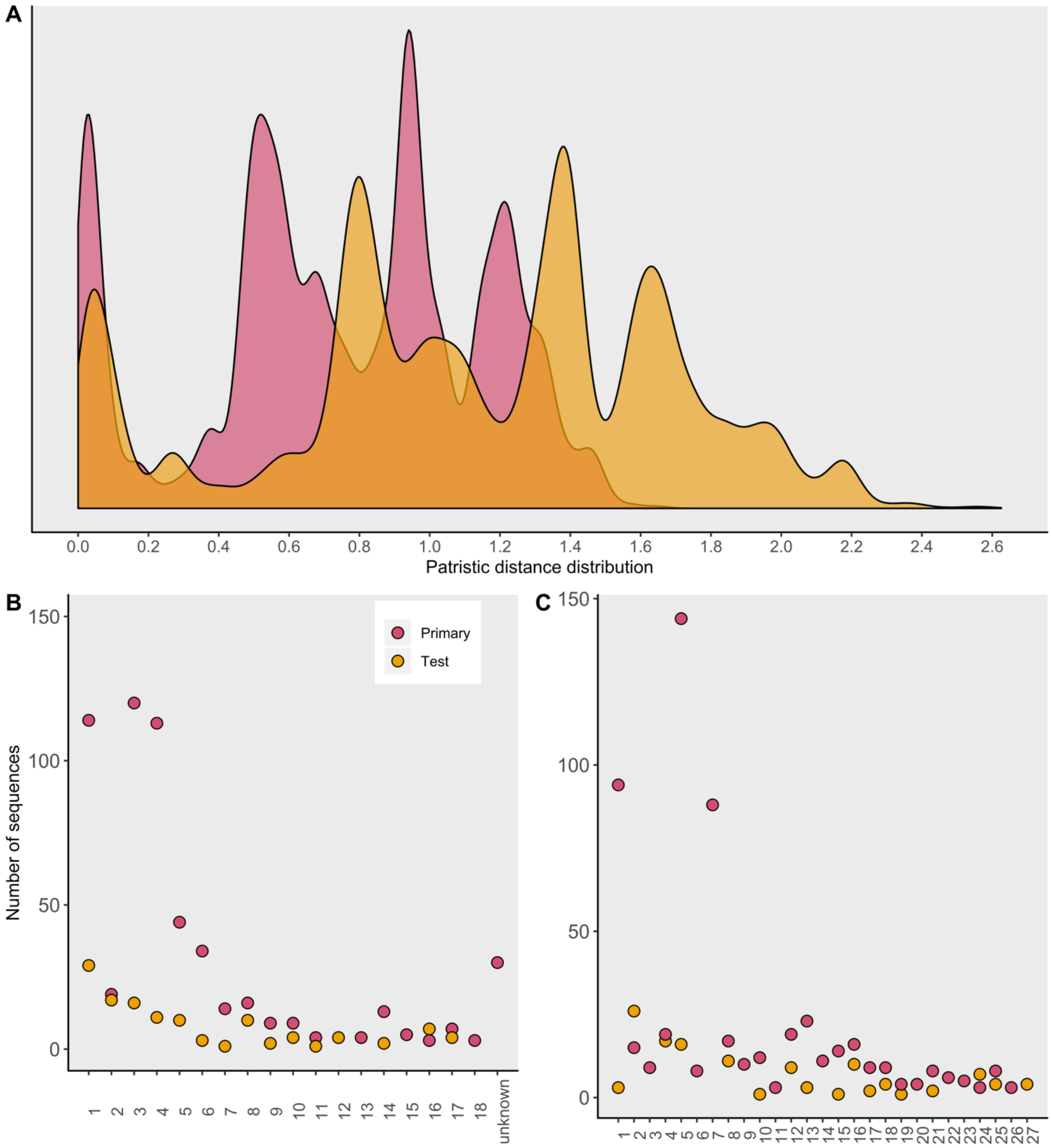


Supplementary Figure 12: A) Density of pairwise patristic distance distribution of the primary and test phylogenies B) Distribution of the number of sequences by existing genotype across the primary and test dataset. C) Distribution of the number of sequences by PhyCLIP genotype across the primary and test dataset. The genotypes predominating in the reference dataset, HPEV-1/PhyCLIP cluster 2, HPEV-3/PhyCLIP cluster 5, HPEV-4/PhyCLIP cluster 8, and HPEV-5/PhyCLIP cluster 12 had the highest frequency of additional viruses. Notably, a disproportionately high number of HPEV-2/PhyCLIP cluster 4 sequences were added to the tree, doubling the representation. Several minor genotypes were not detected in the Malawian cohort.[26]

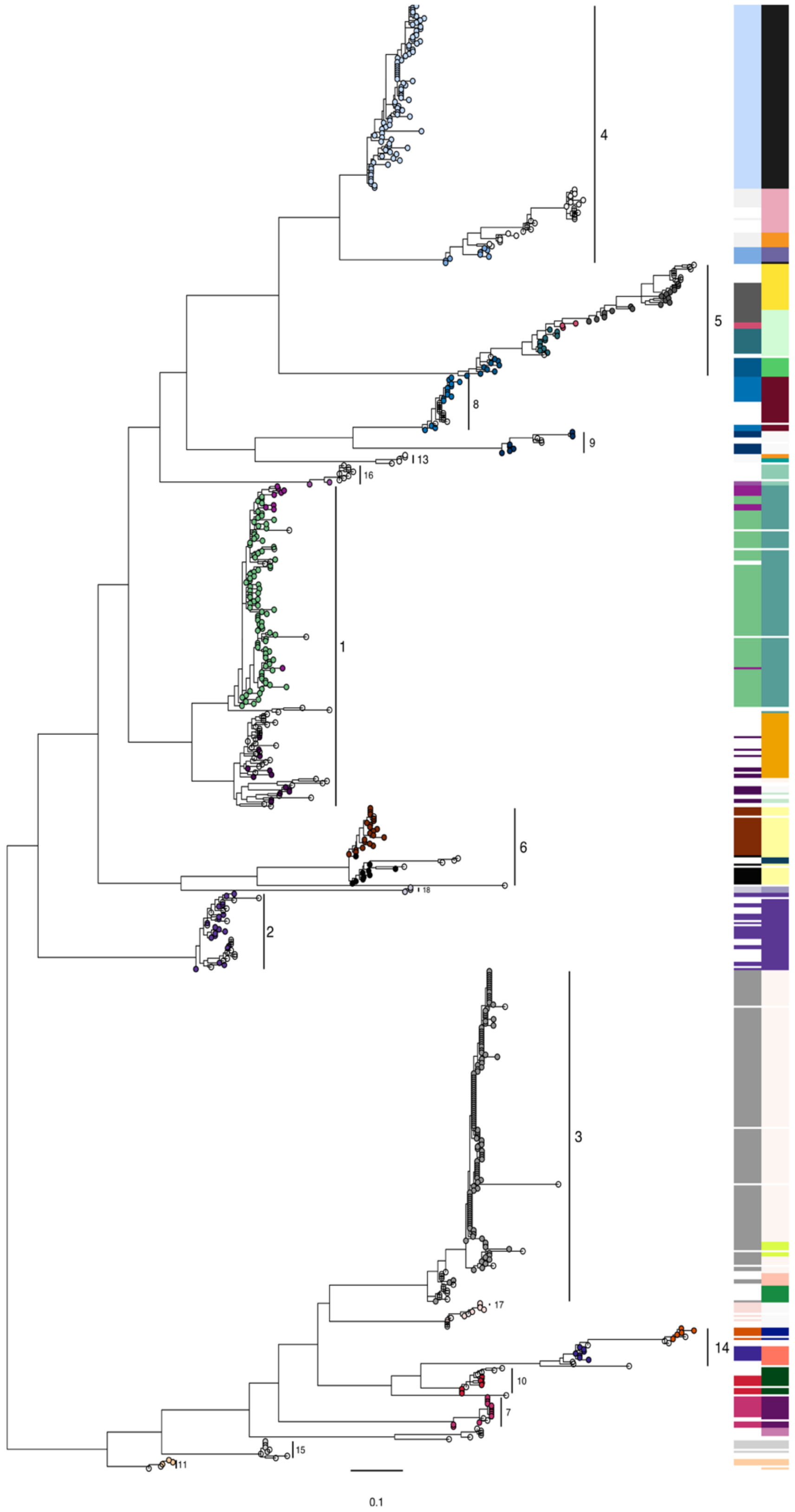


Supplementary figure 13: Reference vs test phylogeny phylogenetic clustering. Annotated by HPeV genotype. Tips coloured by phylogenetic clustering of primary (reference) phylogeny.

### Supplementary tables

Supplementary table 1: Model fit for adaptive branch-site random effects likelihood (aBSREL) branch length re-estimation

| Model | AICc | log L | Parameters |
|---|---|---|---|
| Nucleotide GTR | 132809.28 | -65240.33 | 1161 |
| Baseline MG94xREV | 109680.82 | -52480.25 | 2320 |
| Full adaptive model | 108307.99 | -51584.47 | 2522 |

Baseline MG94xREV refers to the MG94xREV baseline model that infers a single ω rate category per branch. Full adaptive model refers to the adaptive aBSREL model that infers an optimized number of ω rate categories per branch.

Supplementary table 2: aBSREL model complexity for nucleotide phylogeny

| Number of ω rate classes infered | Number of branches with number of rate categories | Percentage of branches with rate categories | Percentage of tree length with the number of rate categories | Number of branches with this number of rate categories |
|---|---|---|---|---|
| **1** | **1052** | **91%** | **0.00%** | **0** |
| **2** | **101** | **8.80%** | **100%** | **0** |

Supplementary table 3: Summary of branches with an uncorrected p-value < 1 in aBSREL full adaptive model

| Branch | Optimized branch length | LRT | Test p-value | Uncorrected p-value | ω distribution over sites |
|---|---|---|---|---|---|
| **Hpev/15/KF626459/Pakistan_2008** | **0.0031** | **15.7784** | **0.1461** | **0.0001** | **$\omega_1$ = 0.00 (100%)<br>$\omega_2$ = 5000 (0.48%)** |
| **MRCA of HPeV 1, 6, 8, 9, 13, 16, 18** | **0.1458** | **15.0254** | **0.2132** | **0.0002** | **$\omega_1$ = 0.00 (96%)<br>$\omega_2$ = 26700 (4.3%)** |
| **MRCA of HPeV 10, 14** | **0.1231** | **14.4636** | **0.2826** | **0.0002** | **$\omega_1$ = 0.230 (96%)<br>$\omega_2$ = 323 (4.5%)** |
| **MRCA 3, 7, 10, 11, 12, 14, 15, 17** | **0.1062** | **14.3319** | **0.3017** | **0.0003** | **$\omega_1$ = 0.00 (92%)<br>$\omega_2$ = 224 (8.1%)** |
| **Hpev/unknown/AB300944/2001** | **0.0248** | **14.2888** | **0.3081** | **0.0003** | **$\omega_1$ = 0.539 (98%)<br>$\omega_2$ = 159 (1.8%)** |
| **MRCA 3, 7, 10, 12, 14, 15, 17** | **0.1363** | **13.4458** | **0.4707** | **0.0004** | **$\omega_1$ = 0.00 (96%)<br>$\omega_2$ = 284 (4.3%)** |
| **Hpev/11/HQ163873/SriLanka/2005** | **0.0254** | **13.2356** | **0.5229** | **0.0005** | **$\omega_1$ = 0.639 (99%)<br>$\omega_2$ = 5250 (0.55%)** |

Supplementary table 4: Mixed Effects Model of Evolution results of episodic diversifying selection at individual sites across HPeV genotypes

| Genotype | Site | LRT | p-value | Number of branches under selection | Total branch length | Nr of sequences |
|---|---|---|---|---|---|---|
| 1 | 137 | 18.29 | 0 | 3 | 16.06 | 117 |
| 1 | 85 | 4.93 | 0.04 | 0 | 7.07 | 117 |
| 1 | 92 | 4.51 | 0.05 | 6 | 4.86 | 117 |
| 2 | 138 | 6.06 | 0.02 | 1 | 4.44 | 37 |
| 3 | 224 | 5.01 | 0.04 | 1 | 3.64 | 120 |
| 4 | 225 | 14.79 | 0 | 4 | 5.95 | 124 |

| | | | | | | |
|---|---|---|---|---|---|---|
| 4 | 229 | 3.98 | 0.06 | 1 | 7.47 | 124 |
| 5 | 222 | 5.49 | 0.03 | 2 | 3.5 | 48 |
| 6 | 84 | 6.07 | 0.02 | 2 | 1.22 | 37 |

Supplementary table 5: Genotype distribution by country

| Country | Number of sequences | Number of genotypes |
|---|---|---|
| Ghana | 112 | 14 |
| India | 83 | 13 |
| Bolivia | 23 | 12 |
| USA | 114 | 6 |
| Netherlands | 276 | 6 |
| Hong Kong | 131 | 6 |
| UK | 38 | 6 |
| Japan | 448 | 5 |
| Pakistan | 15 | 5 |
| Thailand | 10 | 5 |
| Brazil | 9 | 4 |
| Bulgaria | 5 | 4 |
| China | 8 | 4 |
| Finland | 72 | 4 |
| Germany | 11 | 4 |
| Sri Lanka | 15 | 4 |
| Taiwan | 51 | 4 |
| Ivory Coast | 5 | 3 |
| Norway | 23 | 3 |
| South Korea | 27 | 3 |
| France | 37 | 2 |
| Hungary | 8 | 2 |
| Australia | 22 | 1 |
| Bangladesh | 1 | 1 |
| Canada | 1 | 1 |
| Denmark | 1 | 1 |

Supplementary table 6: Spatiotemporal metadata of PeV-A genotypes defined by the current system

| Genotype | Date | Country | Number of sequences |
|---|---|---|---|
| 1 | 1956, 1990-2017 | Brazil, Ghana, Netherlands, UK, USA, Denmark, Finland, India, Japan, Norway, China, Ivory Coast, Hungary, Sri Lanka, Germany, South Korea, Bolivia, Hong Kong, Taiwan, Thailand | 608 |
| 2 | 1956, 2001-3, 2005-9, 2014 | Ghana, Netherlands, USA, India, Germany, UK, Thailand, Japan, Bolivia | 20 |
| 3 | 1990, 1994, 1997, 1999-2016 | Canada, Ghana, Australia, Germany, USA, Norway, Finland, India, France, Netherlands, China, Thailand, Japan, South Korea, Bolivia, Hong Kong, Taiwan, Bulgaria | 614 |
| 4 | 1975, 1979, 1993, 1999, 2001, 2002, 2003, 2004, 2005, 2006, 2007, 2008, 2009, 2010 , 2011, 2012, 2014, 2015, 2016 | Ghana, Netherlands, USA, Thailand, Finland, India, France, UK, Japan, Sri Lanka, Hungary, South Korea, Bolivia, Hong Kong, Taiwan | 115 |
| 5 | 1981-3, 1986, 1992, 2000, 2004-11, 2014-5 | Brazil, Ghana, Netherlands, UK, USA, India, China, Ivory Coast, Hong Kong, Bulgaria | 45 |
| 6 | 2000-9, 2011, 2013-15 | Brazil, Ghana, Netherlands, USA, Norway, Finland, India, China, Germany, UK, Japan, Taiwan, Hong Kong | 43 |
| 7 | 2002, 2003, 2007, 2009 | Ghana, Pakistan, Bolivia, India | 15 |
| 8 | 2006-7, 2010-11 | Brazil, Ghana, India, Bulgaria | 16 |
| 9 | 2002, 2004, 2007 | Ghana, Bolivia | 9 |
| 10 | 2004-8, 2010, 2011 | Ghana, Pakistan, India, Sri Lanka, Bolivia, Bulgaria | 10 |
| 11 | 2004-6 | Sri Lanka, Bolivia, India | 4 |
| 12 | 2002, 2004, 2011 | Pakistan, Bolivia, Bangladesh | 3 |
| 13 | 2005, 2008 | Pakistan, Bolivia | 5 |
| 14 | 2005-7, 2010, 2013, 2016 | Ghana, India, Hong Kong, Thailand | 15 |
| 15 | 2007-8 | Ghana, Pakistan, Bolivia | 10 |
| 16 | 2008 | Bolivia, India | 3 |
| 17 | 2007, 2009, 2014 | Ghana, India, UK, Ivory Coast, China, Ethiopia | 8 |
| 18 | 2007 | Ghana | 3 |

Supplementary table 7: Spatiotemporal metadata of PhyCLIP clusters

| PhyCLIP cluster | Date | Country | Number of sequences | HPeV genotype |
|---|---|---|---|---|
| 1 | 1993-4, 1997-2016 | Ghana, Netherlands, Denmark, Finland, India, Norway, China, UK, Japan, Sri Lanka, Hungary, South Korea, Hong Kong, Taiwan, Thailand | 94 | 1 |
| 2 | 1956, 2004, 2005-11, 2014-15 | USA, India, China, UK, Thailand, Hong Kong | 15 | 1 |
| 3 | 2004, 2007-8, 2010, 2012, 2015 | Ghana, India, Taiwan, Netherlands, Hong Kong | 9 | 1 |
| 4 | 2001-3, 2005-9, 2014 | Ghana, Netherlands, India, Germany, UK, Thailand, Japan, Bolivia | 19 | 2 |
| 5 | 1994, 1997, 1999, 2001-2, 2004-16 | Canada, Ghana, Australia, USA, Norway, Finland, France, Netherlands, China, Japan, South Korea, Hong Kong, Taiwan | 144 | 3 |
| 6 | 2006-09 | Ghana, India | 8 | 4 |
| 7 | 1975, 1979, 1993, 1999, 2001, 2004-12, 2014-16 | Ghana, Netherlands, USA, Thailand, Finland, India, France, UK, Hungary, Sri Lanka, Japan, South Korea, Hong Kong, Taiwan | 88 | 4 |
| 8 | 2002, 2003, 2007 | Ghana, Bolivia | 17 | 4 |
| 9 | 1981-83, 1986, 1992, 2000, 2004-5, 2011 | Netherlands, USA, UK | 10 | 5 |
| 10 | 2007 | Ghana, Netherlands | 12 | 5 |
| 11 | 2004, 2006 | Brazil, Netherlands | 3 | 5 |
| 12 | 2004-11, 2014,-15 | Netherlands, India, China, Ivory Coast, Hong Kong, Bulgaria | 19 | 5 |
| 13 | 2000-1, 2004, 2006-9, 2014-5 | Netherlands, Finland, China, Japan, Taiwan, Hong Kong | 23 | 6 |
| 14 | 2001-02, 2004-08, 2013 | Brazil, Ghana, Netherlands, USA, Finland, India, Japan, Norway | 11 | 6 |
| 15 | 2003, 2007, 2009 | Ghana, Pakistan, Bolivia, India | 14 | 7 |
| 16 | 2006-7, 2010-11 | Brazil, Ghana, India, Bulgaria | 16 | 8 |
| 17 | 2002, 2004, 2007 | Ghana, Bolivia | 9 | 9 |
| 18 | 2004-6, 2008, 2010-11 | Sri Lanka, Pakistan, Bolivia, India, Bulgaria | 9 | 10 |
| 19 | 2004-6 | Sri Lanka, Bolivia, India | 4 | 11 |
| 20 | 2008 | Pakistan | 4 | 13 |
| 21 | 2004, 2007 | Ghana, Netherlands | 8 | 14 |
| 22 | 2005, 2007, 2013, 2016 | India, Hong Kong, Thailand | 6 | 14 |
| 23 | 2008 | Pakistan | 5 | 15 |
| 24 | 2008 | Bolivia, India | 3 | 16 |
| 25 | 2007, 2009, 2014 | Ghana, India, UK, Ivory Coast | 8 | 17 |
| 26 | 2007 | Ghana | 3 | 18 |
